# L-Fucose–Dependent Biofilm Formation by *Escherichia coli* Enhances Polymicrobial Interactions and Antibiotic Tolerance on Urinary Catheters

**DOI:** 10.64898/2026.06.01.729324

**Authors:** Steven M Taddei, Namrata Deka, Adam Marin, Benjamin C Hunt, L Beryl Guterman, Min Ma, Jun Qu, Chelsie E. Armbruster

**Affiliations:** Department of Microbiology and Immunology, Jacobs School of Medicine and Biomedical Sciences, State University of New York at Buffalo, Buffalo, NY, USA; Department of Pharmaceutical Sciences, School of Pharmacy and Pharmaceutical Sciences, State University of New York at Buffalo, Buffalo, NY, USA; NYS Center of Excellence in Bioinformatics and Life Sciences, Buffalo, NY, USA

**Keywords:** Polymicrobial, biofilm, urinary tract, fucose, E. coli, catheter, P. mirabilis, E. faecalis

## Abstract

Urinary tract infections are common healthcare associated infections, a large subset of which are caused by indwelling catheters. Long term catheterization causes persistent, asymptomatic, polymicrobial colonization despite catheters changes and antibiotic usage. In these polymicrobial populations, *P. mirabilis*, *E. faecalis*, and *E. coli* were found as the most common co-colonizing species. We investigated how interactions between *P. mirabilis*, *E. coli*, and *E. faecalis* contribute to biofilm formation and colonization of urinary catheters. Our results show that the interaction between these three species leads to enhanced biofilm biomass driven by an increase in total protein content of the biofilm. Biofilm enhancement required all three species and was also media-dependent, especially for dual-species combinations. Importantly, triple species biofilms also demonstrate biofilm enhancement when established under flow conditions in a biofilm reactor model using silicone urinary catheters. Additionally, triple species biofilm enhancement occurred in co-colonizing isolates from catheterized patients and was found to be specific to interactions between these three species. Triple species biofilms also demonstrated a species-dependent resistance to two commonly used antibiotics, ciprofloxacin and nitrofurantoin. By examining priority effects, *E. coli* was found to be the main facilitator of biofilm enhancement in a flow model. Finally, proteomics revealed that an L-fucose utilization pathway in *E. coli* was a key contributor to triple species biofilm enhancement. Overall, our results demonstrate the significant impact of polymicrobial interactions on biofilm formation in the catheterized environment and highlight ways in which complex microbial interplay and priority effects can shape the establishment of persistent colonization.

## Introduction/Background

Urinary tract infections (UTI) are commonly-experienced infections worldwide that affect 40-60% of women and 12-15% of men (1). These infections are often considered to be monomicrobial and most frequently attributed to *Escherichia coli*, a gram-negative bacterium (2, 3). Up to 80% of healthcare-associated UTIs are due to the presence of an indwelling catheter, resulting in catheter associated urinary tract infection (CAUTI). CAUTIs are especially common in vulnerable populations such as the immunocompromised and residents in nurse homes, and these infections are often recurrent and associated with an increased incidence of urolithiasis, renal damage, and secondary bacteremia (4–7). CAUTIs also tend to be caused by a more diverse range of microbes than UTIs, and are often polymicrobial (7–9). For example, one study of clinically-diagnosed and treated CAUTIs reported that out of 182 cases, 57 (31%) had at least two species reported (10). In dual species cultures, *Enterococcus* spp. was found the most frequently in 23 samples (40%), followed by *Proteus mirabilis* (20, 35%), *Pseudomonas* aeruginosa (16, 28%), and *Escherichia coli* (14, 25%).

The catheterized bladder provides a unique niche environment that promotes the colonization and growth of uropathogens, thereby facilitating polymicrobial colonization and subsequent infection. Urinary catheters obstruct the natural flushing of the urethra and the complete voiding of urine from the bladder, allowing for a reservoir of urine below the drainage eyelet of the catheter. Urine contains metabolites that support bacterial growth, such as amino acids, organic acids, and iron. Additionally, insertion of the catheter can lead to uroepithelial cell damage, which recruits damage-response and inflammatory proteins that contribute to bacterial cell adhesion (11–13). These factors facilitate microbial colonization and proliferation in the urinary tract, as well as microbial biofilm formation. Catheter biofilms are typically polymicrobialand provide protection from the host immune system and antibiotics, facilitating persistence (14, 15). Due to these factors, bacteria are frequently present in urine specimens from individuals with long-term catheters in the absence of infection signs or symptoms, which is considered to be catheter-associated asymptomatic bacteriuria (CA-ASB).

CA-ASB is common and found in the vast majority of patients after a month of indwelling catheter use, and is often polymicrobial (7, 16). Two recent studies have examined the microbial composition of CA-ASB in different patient populations with long-term catheters: nursing home residents, and community dwelling individuals (7, 9). In the nursing home setting, 226 out of 233 urine samples (97%) were found to be polymicrobial, averaging ∼4 species per sample. *Enterococcus faecalis* (70%), *Proteus mirabilis* (51%), and *Escherichia coli* (38%) were the top species isolated from these samples, with all three of these microbes being found together in 20% of the samples(7). Similar findings were observed in community-dwelling patients; 290 out of 363 (80%) samples were polymicrobial, averaging ∼3 species present per sample. *E. faecalis* (51%), *P. mirabilis* (29%), and *E. coli* (20%) were also among the top species isolated and found co-colonized in 15% of the samples (9). These studies suggest that long-term catheterization can lead to stable polymicrobial colonization, and the persistent co-colonization of *P. mirabilis*, *E. coli*, and *E. faecalis* suggests a possible synergistic partnership between these species.

Each bacterium has an arsenal of virulence and fitness factors that assist in colonization and pathogenesis, many of which are also likely to be important contributors to polymicrobial infection. *E. faecalis* is a gram-positive bacterium that can colonize diverse environments and is associated with wound infections, bloodstream infections, and CAUTI (17). UTI-related *E. faecalis* strains harbor notable virulence factors that manipulate host factors to increase colonization, such as immune modulation through suppressing TLR signaling (18–20) and increased binding to the catheterized bladder environment by sequestering host factors like fibrinogen (12, 21). *E.coli* is a gram-negative bacterium responsible for more than 50% of all UTIs (22). Uropathogenic *E. coli* strains are characterized by an increased ability to cause infection outside of the intestinal tract, namely in the urinary tract, bloodstream, and cerebrospinal fluid (3). Key pathogenic factors of *E. coli* include the ability to rapidly grow in urine via flexible metabolism and numerous iron acquisition systems, including secreted siderophores that can be pirated by other bacteria (23). *P. mirabilis* is a gram-negative bacterium known for its urease activity, which increases urine pH and causes the formation of bladder, kidney stones, and crystalline biofilms that facilitate bacterial adherence and cause tissue damage. CAUTIs caused by *P. mirabilis* are associated with high mortality rates due to bacteremia and sepsis following inf*ec*tion, further increased during polymicrobial infection (24).

During polymicrobial colonization, uropathogens must either compete or cooperate with co-colonizing partners to persist. Common themes that have arisen from studies of polymicrobial interactions between uropathogens have demonstrated that both cooperation and competition can impact viability in one or both bacteria, adhesion and biofilm formation, tissue damage and disease progression, and modulation of host immune responses (11, 23, 25–27). For example, when *E. coli* and *E. faecalis* are co-inoculated in iron-limited environments, an increase in biofilm formation and *E. coli* survival was observed and this phenotype was linked to the secretion of excess ornithine by *E. faecalis* (23). Cooperation between *E. coli* and *E. faecalis* was further observed in a longitudinal study investigating microbial co-occurrences in catheterized patients. In this context, co-culture of *E. coli* and *E. faecalis* in artificial urine media increased *E. faecalis* viability compared to single species cultures (9). Our laboratory recently demonstrated that polymicrobial interactions between *P. mirabilis* and *E. faecalis* can also be beneficial to both microbes, resulting in increased biofilm biomass and infection severity without altering viability of either species, and that metabolic interplay involving ornithine secretion and arginine biosynthesis contributes to this process (11, 26). *P. mirabilis* urease activity is also increased during *in vitro* co-culture with either *E. coli* or *E. faecalis* in human urine, and the resulting pH increase can decrease bacterial viability *in vitro* (11). However, *P. mirabilis* coinfection in a mouse model of CAUTI increased *E. coli* CFUs in the bladder and kidneys (28).

Notably, most polymicrobial studies involving uropathogens have been performed in standard laboratory media, and the interactions between species may differ when examined in more physiologically relevant media such as human urine. Additionally, most polymicrobial studies involving uropathogens focus on interactions between only two bacterial species, as described above. Thus, there is a gap in our understanding of the mechanisms that contribute to successive colonization and persistence of 3+ species during long-term catheterization and how interactions between more complex polymicrobial communities influence infection risk and severity. In this study, we investigated how triple species interactions between *P. mirabilis*, *E. coli*, and *E. faecalis* contribute to biofilm formation and colonization of the urinary catheter environment. We found that co-culture of *P. mirabilis*, *E. coli*, and *E. faecalis* increases biofilm formation through a specific increase in total protein within the biofilm. Biofilm enhancement occurred in clinical co-isolates as well as commonly-used strains, and represents a unique relationship between these three uropathogens. Furthermore, *E. coli* was found to be the main driver of the enhanced biofilm biomass phenotype, and L-fucose metabolism was a key contributor to initial *E. coli* colonization of the catheterized environment and subsequent interaction with *P. mirabilis* and *E. faecalis*. Altogether, this work broadens our understanding of the CA-ASB environment and facilitates future studies aimed at examining the contribution of synergistic polymicrobial interactions in the development of unique catheter biomes as well as their contribution to onset of symptomatic infection. Ultimately, continued studies may reveal processes that can be targeted to decrease the incidence and severity of CAUTIs in patients with long-term catheters.

## Results

### Polymicrobial interactions during planktonic growth can decease viability in a urease-dependent manner

To determine whether interactions between *P. mirabilis, E. coli,* and *E. faecalis* exhibit competitive or cooperative behaviors during planktonic growth, colony forming units (CFUs) of each species from single, double, and triple cultures were determined over a time-course in tryptic soy broth with glucose (TSBG), artificial urine media (AUM), and human urine (Figure 1). We hypothesized that viability would be unaltered in nutrient rich TSBG while in AUM and human urine there may be competition for nutrients that could decrease CFUs of one or more species. *E. faecalis* and *E. coli* also secrete factors that can increase *P. mirabilis* urease activity (28), so we further hypothesized that the viability of one or more species could be decreased in a urease-dependent manner if they do not tolerate the rapid rise in pH.

**Figure 1:**
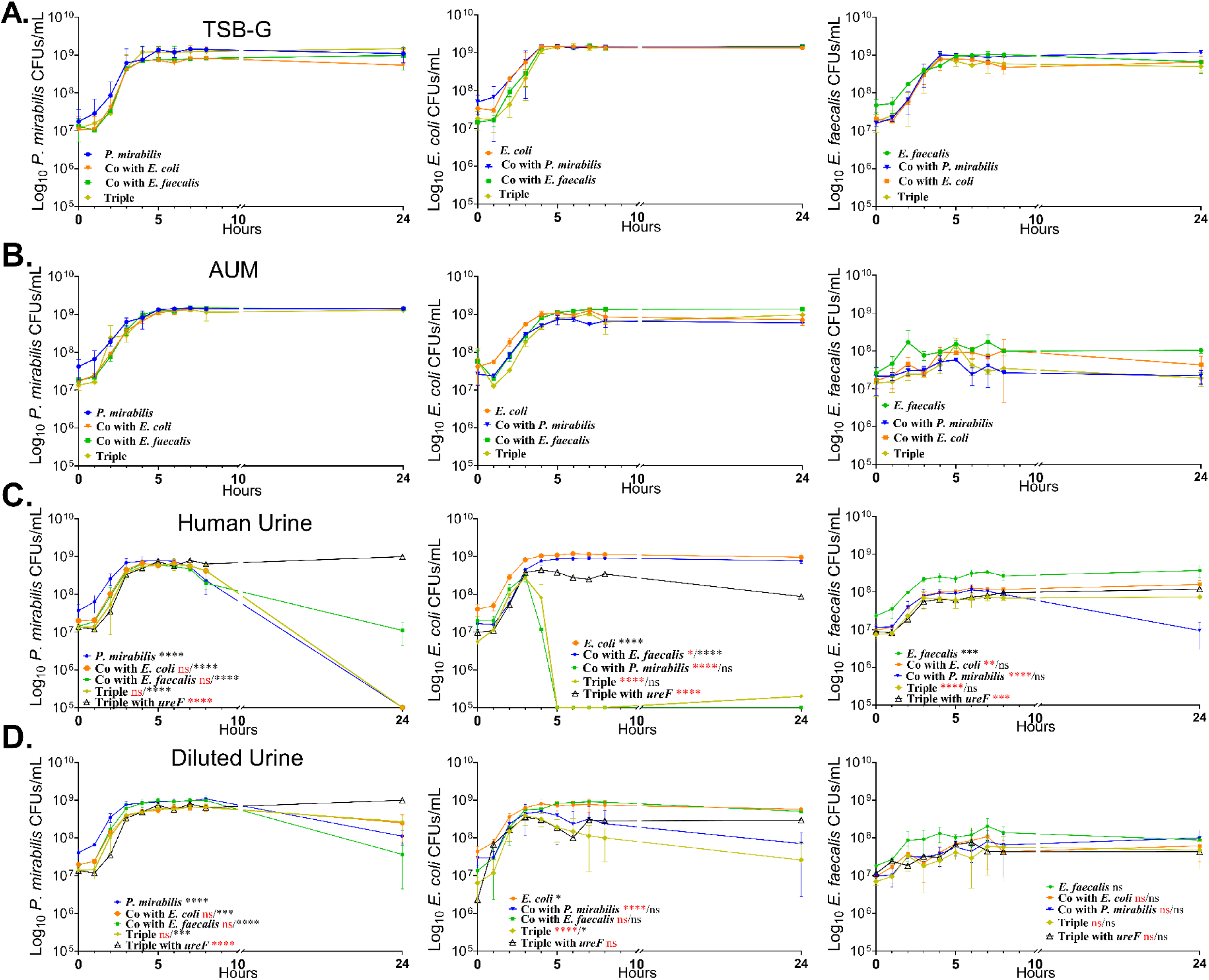
Polymicrobial growth curves suggest a urease-mediated decrease in cell viability, which is not due to competition. *P. mirabilis*, *E. coli*, *E. faecalis,* and a *P. mirabilis* urease mutant (*ΔureF*) were cultured in single, double, and triple species combinations in (A) TSB-G, (B) artificial urine medium (AUM), (C) human urine, and (D) human urine diluted with 0.9% saline to a specific gravity of 1.004. Individual species growth was assessed by differential plating hourly from 0-8h and at 24 hours to determine CFU/mL. Data shows the mean ± standard deviation for at least 2 independent experiments with 3 replicates each. Statistical comparisons were made to the *ureF*-triple group (black, asterisk) or to single species (red, asterisk) by one-way ANOVA, using a multiple comparison test; *, P<0.05; **, <0.01; ***, P<0.001; ****, P<0.000

Single species cultures exhibited robust growth in all media types, although *E. faecalis* only exhibited a modest overall increase in viability when cultured in either AUM or human urine. Growth of all three species in either TSBG or AUM remained consistent between single and polymicrobial cultures (Figure. 1A-B), demonstrating that there is no obvious competition or cooperation between these species that impacts viability. However, all three species exhibited reduced viability in human urine cultures containing *P. mirabilis* (Figure. 1C). To test whether decreased cell viability was urease-mediated, co-cultures in urine were repeated using a *P. mirabilis* urease mutant (*ureF,* Figure. 1C). Loss of *P. mirabilis* urease activity largely restored viability of all three species, suggesting that the decrease in cell viability in human urine was driven by a urease-dependent increase in pH rather than direct competition between species.

To investigate whether decreasing the concentration of urea and other metabolites could push the bacteria towards competitive interactions, growth curves were repeated using human urine diluted ∼1:1 with 0.9% saline to achieve a specific gravity of 1.004, which we previously found to slow down urease-mediate pH changes for *P. mirabilis* (29). All three species exhibited robust growth in dilute human urine, and no significant differences in viability were observed between culture conditions (Figure. 1D). Thus, the urease-dependent decrease in viability that occurs in human urine is tempered by diluting the urine and reducing the urea concentration, but the resulting reduction in nutrient abundance does not force any observable competition between species.

### Polymicrobial interactions enhance biofilm biomass in a media-dependent manner

To understand the impact of polymicrobial interactions on biofilm development, single (Pm, Ec, Ef), double (Pm/Ec, Pm/Ef, Ec/Ef), or triple (Pm/Ec/Ef) species biofilms were established in tissue culture treated 24 well plates in either TSBG, AUM, human urine, or dilute human urine, and biofilm biomass was assessed using crystal violet staining after a 24 hour stationary incubation (Figure 2). Crystal violet readings were normalized to single species *E. faecalis* biofilms within each experiment to facilitate comparisons across independent experiments, and polymicrobial biofilms were only considered to exhibit enhanced biomass if the crystal violet staining was significantly greater than the single species biofilms formed by each species in the mixture. The biomass of single species biofilms varied across species, with *E. faecalis* forming the most robust biofilms in urine and dilute urine, *P. mirabilis* forming the most robust biofilms in AUM, and *E. coli* generally exhibiting lower biomass than the other species except when cultured in undiluted urine. Co-culture of *P. mirabilis* with *E. coli* did not significantly alter biofilm biomass in any of the media types, but a bimodal distribution was observed for biofilms formed in human urine (Figure. 2C). Co-culture of *P. mirabilis* with *E. faecalis* resulted in an increase in biofilm biomass in all media types, consistent with our prior data (26). Co-culture of *E. coli* with *E. faecalis* did not alter biofilm biomass in most media types, but robust enhancement was observed in AUM (Figure 2B). In contrast, triple species biofilms always exhibited enhanced biomass, suggesting that the interactions driving dual-species enhancement are media-dependent while those driving triple-species enhancement are independent of media type (Figure 2A-D).

**Figure 2:**
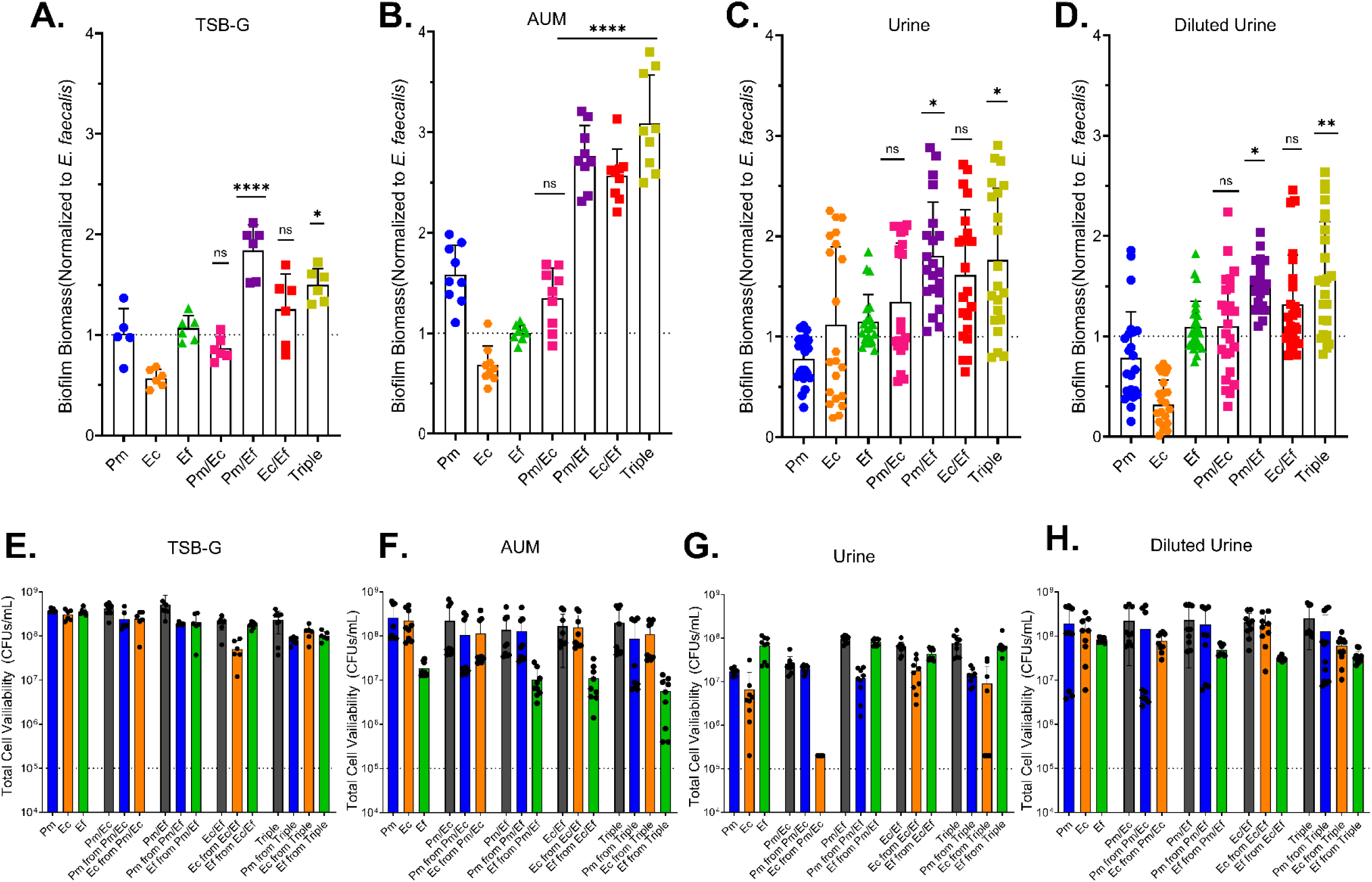
Triple species biofilms exhibit consistent biomass enhancement, while dual species biofilm enhancement is variable and media-dependent. Single (blue, orange, green), double (fuchsia, purple, red), and triple (yellow) species combinations were grown in 24 well plates at 37°C for 24h in (A,E) TSB-G, (B,F) AUM, (C, G) human urine, or (D, H) diluted urine. (A-D) Biofilm biomass was assessed by crystal violet staining read at an absorbance of 570 nm. Polymicrobial combinations were compared to the highest biofilm-forming single species constituent. Error bars represent mean and SDs from at least 2 independent experiments with at least 3 biological replicates each. Stats were determined by Kruskal-Wallis test with multiple comparisons; *, P<0.05; **, <0.01; ****, P<0.001; ns, non-significant. (E-H) Total (grey) and individual species (blue, orange, green) cell viability was determined by differentially plating in single, double, and triple species combinations in (E)TSB-G, (F) AUM, (G) human urine, or (H) dilute urine from 24-hour biofilms.

To determine if cell viability was altered in different media types, differential plating was first used to quantify each species within the single, double, and triple species biofilms to determine whether increased biofilm biomass is due to an increase in abundance of any one species within the biofilm (Figure 2E-G). Total CFUs were largely consistent across all combinations and media types, with no significant differences observed for double or triple species biofilms compared to single species biofilms. Similar trends were observed when examining the CFUs of each individual species, although *E. coli* exhibited a striking decrease in viability when co-cultured with *P. mirabilis* in urine (Figure 2E). This is likely the result of urease-mediated pH changes, as in Figure 1C. Thus, cell viability remains unaltered in polymicrobial biofilms and is not contributing to biofilm enhancement.

### Triple species biofilm biomass enhancement occurs with clinical isolates of P. mirabilis, ***E. coli, and E. faecalis but not with other species***

Considering that *P. mirabilis, E. coli,* and *E. faecalis* were recently observed to persistently co-colonize for several months in patients with indwelling urinary catheters (7, 9) clinical co-isolates of each species were used to determine whether polymicrobial biofilm enhancement also occurs in isolates that have co-adapted within the urinary tract. To do this, isolates were chosen from three study participants: one who was co-colonized by *P. mirabilis* and *E. faecalis* for three weeks and then acquired *E. coli* (Figure 3A), one who was co-colonized by *E. faecalis* and *E. coli* for fifteen weeks and then acquired *P. mirabilis* (Figure 3B), and one who was co-colonized by all three species at baseline and maintained co-colonization for the duration of the study (Figure 3C). Single, double, and triple species biofilms were established in diluted human urine and biofilm biomass was measured by crystal violet staining as previously described. All triple species biofilms exhibited a significant increase in biofilm biomass compared to the single species biofilms. All *Pm/Ef* co-cultures also exhibited an increase in biofilm biomass in all three participant sets, while *Pm/Ec* and *Ec/Ef* co-cultures exhibited greater variability and significant enhancement was only observed for isolates from 2 of the 3 participants. These data suggest that the interactions that drive enhancement between these three species are common even in co-colonizing isolates and could contribute to catheter colonization and persistence.

**Figure 3:**
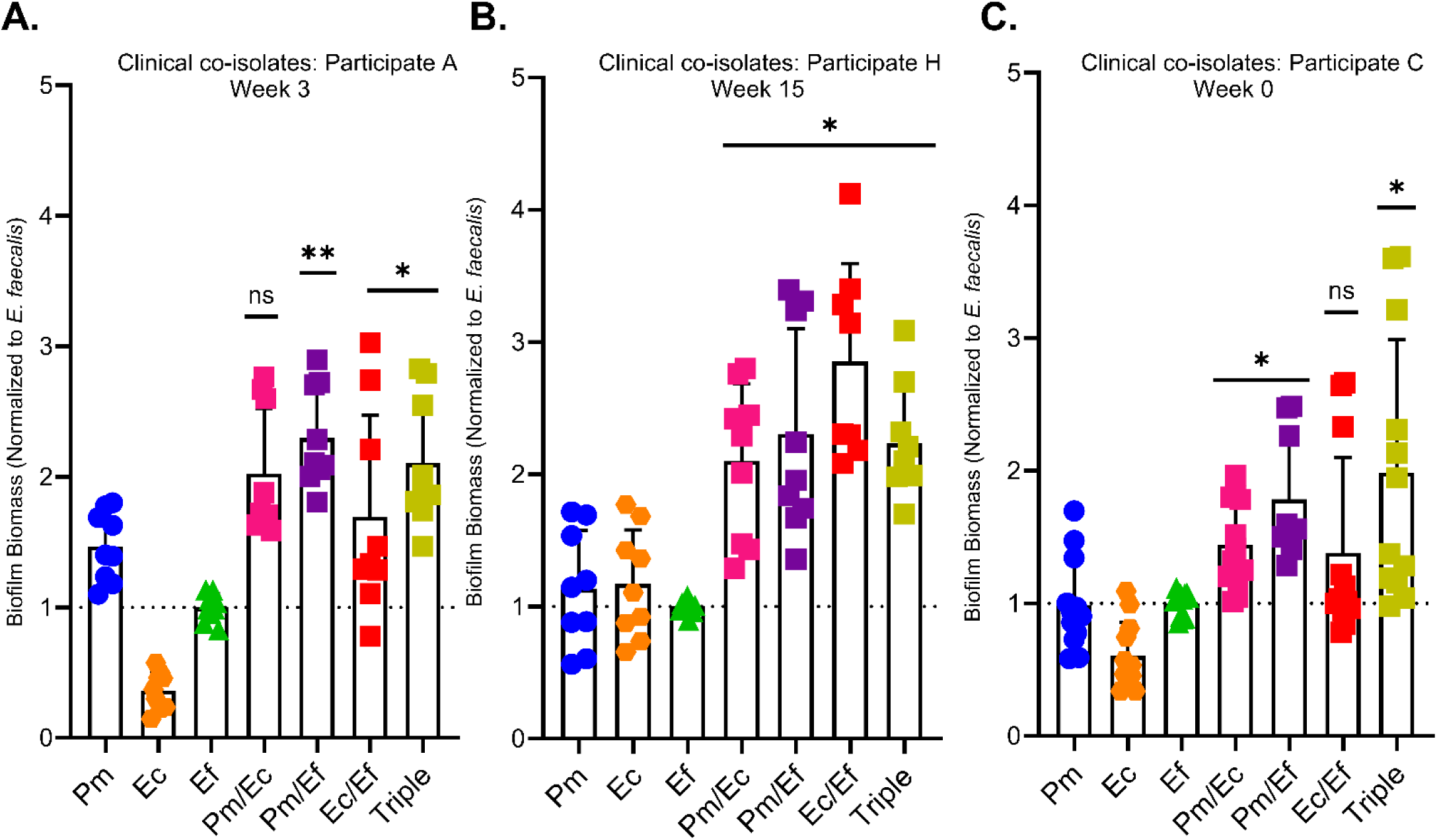
Triple species biofilms formed by clinical co-isolates also exhibit enhanced biomass. Single, double, and triple species biofilms were formed in diluted urine with co-isolated species from (A) participant A, week 3, (B) participant H, week 15 and (C) participant C, week 0. Biofilm biomass was determined by crystal violet staining after 24 hours in diluted urine. Error bars represent mean and SDs 3 independent experiments with 3 biological replicates each. Stats were determined by comparison to the highest single species biofilms using a Kruskal-Wallis test with multiple comparisons; *, P<0.05; **, <0.01; ns, non-significant.

To determine whether triple species biofilm enhancement is specific to the interactions between *P. mirabilis, E. coli,* and *E. faecalis,* additional biofilms were established by substituting out one of these species for other common co-colonizing uropathogens: *Morganella morganii, Providencia stuartii,* or *Staphylococcus aureus* (Supplemental Figure 1). Interestingly, biomass enhancement was not observed for any of the biofilms in which *P. mirabilis, E. coli,* or *E. faecalis* had been swapped out for a different species (Supplemental 1A). These data suggest that a unique interaction occurs when *P. mirabilis*, *E. coli*, and *E. faecalis* co-colonize, leading to enhanced polymicrobial biofilm biomass.

### Triple species interactions increase biofilm biomass on silicone surfaces and under flow conditions

Foley catheters are standard medical devices used to assist in various instances of urinary incontinence and retention. These medical devices are constructed from either latex or silicone; use of latex catheters is no longer as common due to the frequency of patient allergies and the promotion of microbial colonization due to the pitted nature of the material (30–32). Silicone catheters are considered to be more hypoallergenic, cause less irritation, and have a lower rate of encrustation (33). Despite this reasoning, CAUTIs still occur with silicone material, which suggests that microbial attachment to silicone surfaces is pivotal in biofilm formation and proliferation. To determine if polymicrobial interactions lead to increased biofilm biomass on silicone surfaces, 8 mm sterile silicone discs were fitted into untreated 24 well plates containing dilute human urine and inoculated as described above. The discs were incubated for 24 hours, gently washed, and stained with crystal violet to quantify biofilm formation. In contrast to biofilms formed in tissue culture treated 24 well plates, none of the dual species biofilms exhibited enhanced biomass on the silicone discs (Figure 4A). However, the triple species biofilms still demonstrated enhanced biomass compared to all single species biofilms, indicating that interactions between these species can facilitate biofilm formation on silicone surfaces.

**Figure 4:**
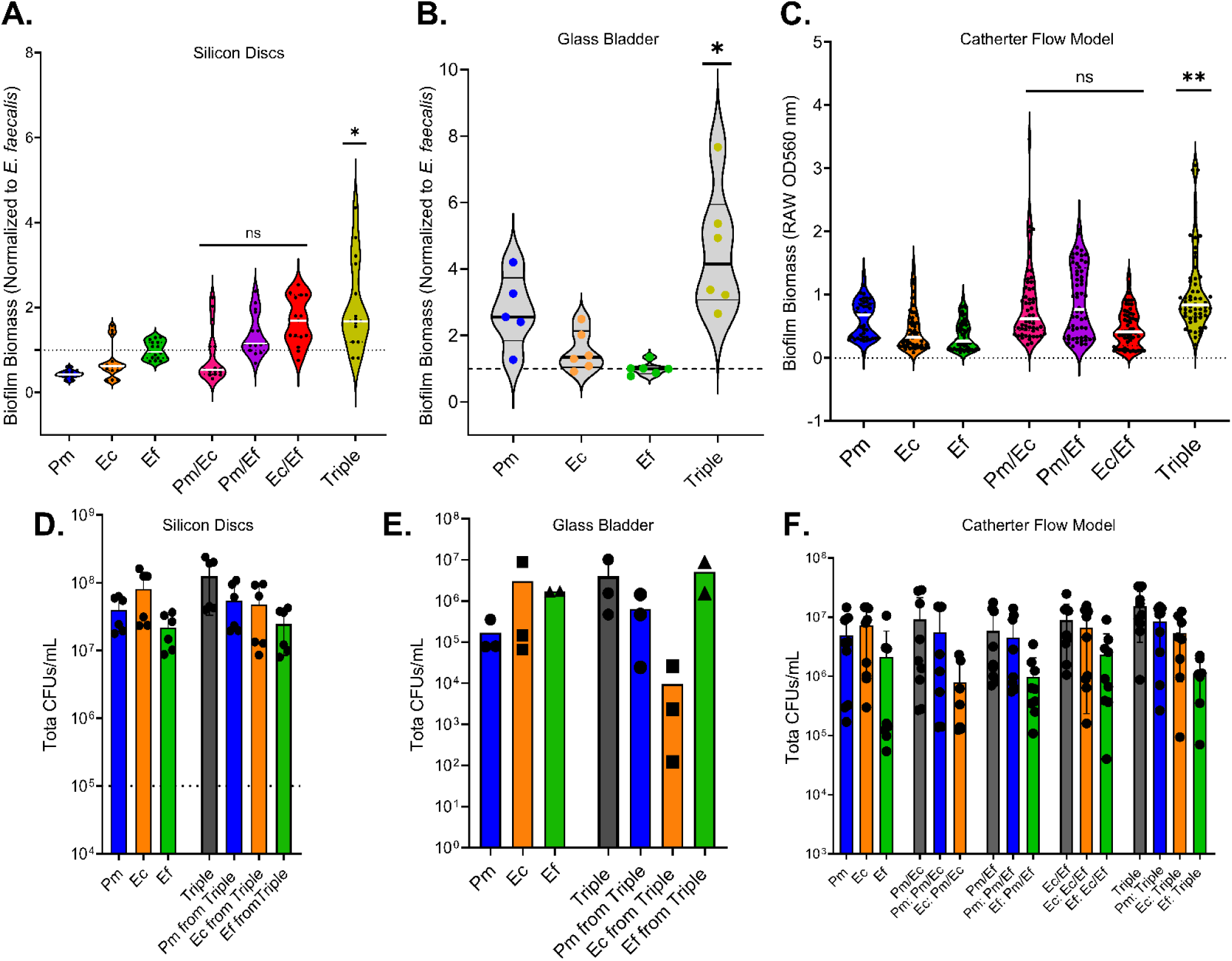
Triple species biofilms still exhibit enhanced biomass when formed on silicone surfaces and under flow conditions. (A) Single, double, and triple species biofilms formed on silicone discs in diluted urine. (B) Single and triple species biofilms formed in a glass bladder model with diluted urine flow. (C) Single, double, and triple species biofilms formed in a catheter biofilm reactor with diluted urine flow. Biofilm biomass was assessed using crystal violet and read at OD560. Data shows at least 2 biological replicates from 2 independent experiments. Statistical significance was determined by ordinary one-way ANOVA with multiple comparisons test or by Kruskal-Wallis with multiple comparisons; *, P<0.05; **, <0.01; ns, non-significant. Total viability and CFU/mL of each individual species were also assessed for (D) silicone disc biofilms, (E) catheter biofilms in the glass bladder model, and (F) catheter biofilms in the periodic flow model. CFUs were determined by differential plating from at least 2 independent experiments with at least two replicates.

Biofilms that form within a urinary catheter will be exposed to liquid flow rather than remaining stationary. To next determine whether enhancement occurs under flow conditions, triple species biofilms were established in a glass bladder model system as previously described (26). This model consists of a 500-milliliter double walled glass vessel in which a Foley catheter is placed and the balloon inflated to form a seal, and human urine is introduced at a constant rate via a peristaltic pump. Due to the large volume of pooled human urine required for these experiments, only single and triple species biofilms were assessed. However, the triple species inocula again resulted in enhanced biomass (Figure 4B), confirming that the polymicrobial interactions driving this phenotype still occur in the catheter lumen under flow conditions.

To further examine biofilm formation during catheterization, including for dual species combinations, a modified biofilm reactor system was developed to reduce the amount of human urine needed for testing (Supplemental Figure 2). The eyelet, balloon, and sampling port were aseptically removed from a silicone Foley catheter, and the remaining length of catheter tubing was placed in a reaction chamber to be maintained at 37°C and attached to a peristaltic pump that flushes 2 ml of urine through the catheter lumen once per hour. Importantly, sterile human urine is left within each catheter for 24 hours before inoculation to allow for development of a conditioning film to promote bacterial attachment. Each inoculum was allowed to incubate within the catheter lumen 1 hour before initiating urine flow from the peristaltic pump. After 24 hour incubation, total biofilm biomass was assessed by staining the catheter lumen with crystal violet and sectioning catheters into 10 mm segments. Variability in staining was observed across the catheter length for all inocula and was most pronounced for the polymicrobial biofilms (Supplemental Figure 2A). None of the dual species biofilms exhibited a statistically significant increase in biomass, although the triple species biofilm exhibited clear enhancement throughout the length of the catheter (Figure 4C).

To again determine whether biofilm enhancement was a reflection of bacterial viability in these models, CFUs were quantified for each silicone surface experiment. For all experiments, viability remained uniform between single species and polymicrobial biofilms (Figure 4D-G) except for a modest decrease in *E. coli* CFUs during triple species infection in the glass bladder model (Figure 4E). Taken together, these data indicate that biofilm enhancement is likely due to a specific interaction that mediates biofilm development rather than promotes cell viability.

### Inoculation order influences triple species biofilm enhancement under flow conditions

To determine if initial colonization order impacts biomass enhancement, catheters in the modified biofilm reactor system were inoculated with each species alone or dual species combinations at time 0, the inocula were allowed to establish for 1.5 hours, and then the remaining species were introduced (Figure 5A). The impact of priority effects was assessed by comparing to single species biofilms as well as a triple species condition in which all three species were simultaneously inoculated, as done previously. When *E. coli* was included in the initial inoculum, final biofilm biomass was enhanced to a similar level as when all three species were inoculated together at time 0. In contrast, when *P. mirabilis* or *E. faecalis* colonized first, either alone or together but in the absence of *E. coli*, there was a significant decrease in final biofilm biomass compared to when all three species were inoculated together at time 0. Interestingly, inoculation with dual species combinations and then adding the third species resulted in the same overall biomass as the triple species inoculum at time 0, although the median biomass value was the lowest when *E. coli* was missing from the initial inoculum (Figure 5A). Taken together, these data suggest that *E. coli* may be the key contributor to facilitating biomass enhancement.

**Figure 5:**
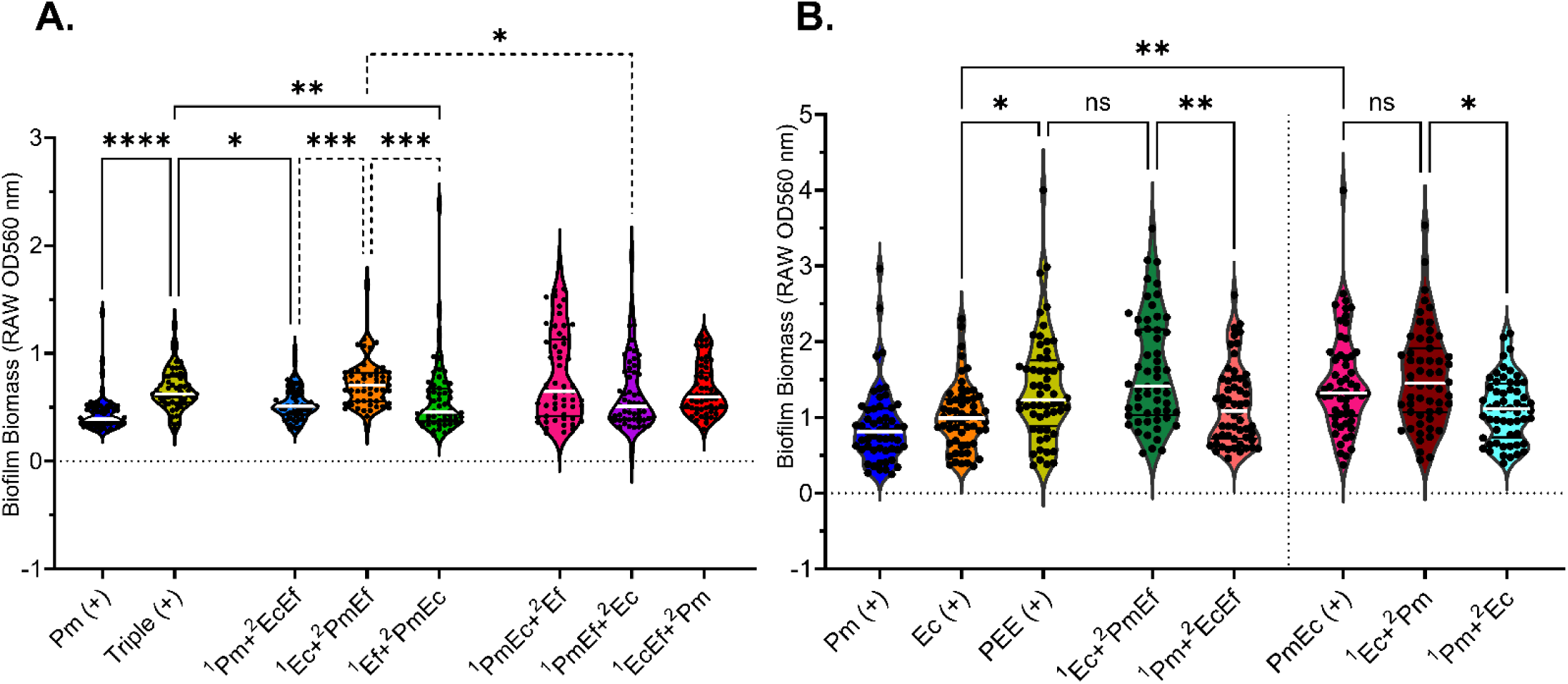
Initial colonization by *E. coli* modulates the catheter environment to facilitate polymicrobial biofilm enhancement. Priority effect experiments were performed under flow conditions in which (A) inoculations were 1.5 hours apart and biomass assessed at 24 hours or (H) inoculations were 24 hours apart and biomass assessed at 48 hours. Pm (blue,+), Ec (orange,+), or Triple (yellow,+) were seeded at time 0 for comparison, and numbers (1,2) indicate which species or combinations were seeded first and which were introduced after the first incubation. Biofilm biomass was assessed by crystal violet staining. Data shows at least 1 biological replicate from 4 independent experiments per condition. Statistical significance was determined by Kruskal-Wallis with multiple comparisons; *, P<0.05; **, P<0.01; ***, P<0.001; ****, P<0.0001; ns, non-significant

To further examine *E. coli* as a key contributor to biofilm enhancement, a 48 hour priority experiment was performed in which the initial inocula were established for 24 hours before the next inoculum was introduced (Figure 5B). The polymicrobial biofilms were then incubated for an additional 24 hours, and biofilm biomass was assessed as previously described. Inoculation with all three species at time 0 resulted in a significant increase in 48 hour biofilm biomass compared to inoculation with either *P. mirabilis* or *E. coli* alone (Figure 5B). Biofilms that contained *E. coli* at time 0 exhibited a significant enhancement compared to the single species biofilms and rivaled the triple species control. However, biofilm biomass was significantly reduced when the initial inoculum contained *P. mirabilis* alone. Overall, these data strongly suggest that *E. coli* is modulating the catheter environment in a way that promotes triple species biofilm development.

### Polymicrobial biofilms have a species-specific impact on antibiotic susceptibility

Biofilms are known to provide protection from antibiotics and are thought to play a role in recurrent UTI and CAUTI infections. Urinary tract infections are commonly treated with ciprofloxacin and nitrofurantoin, antibiotics that inhibit bacterial growth by disrupting DNA replication and protein synthesis, respectively (34–40). To determine whether polymicrobial biofilm formation decreases efficacy of common antibiotics, single and triple species biofilms were formed in diluted human urine for 24 hours in 24 well plates, supplied with fresh urine containing increasing concentrations of either ciprofloxacin or nitrofurantoin, and then incubated for an additional 24 hours. Biofilm CFUs were then quantified to assess species viability after exposure. Single species *P. mirabilis* biofilms were sensitive to concentrations of ciprofloxacin >1 ug/mL, and triple species biofilms provided no significant protection for *P. mirabilis* (Figure 6A). However, viable *P. mirabilis* was recovered from a greater number of polymicrobial catheter segments after treatment with 5 ug/mL ciprofloxacin than from single species segments, suggesting a potentially limited amount of protection provided by the polymicrobial biofilm. In contrast, *P. mirabilis* biofilms were largely resistant to nitrofurantoin at all concentrations, although several of the replicates exhibited cell death at high concentrations of (200 and 250 ug/mL, Figure 6D).

**Figure 6:**
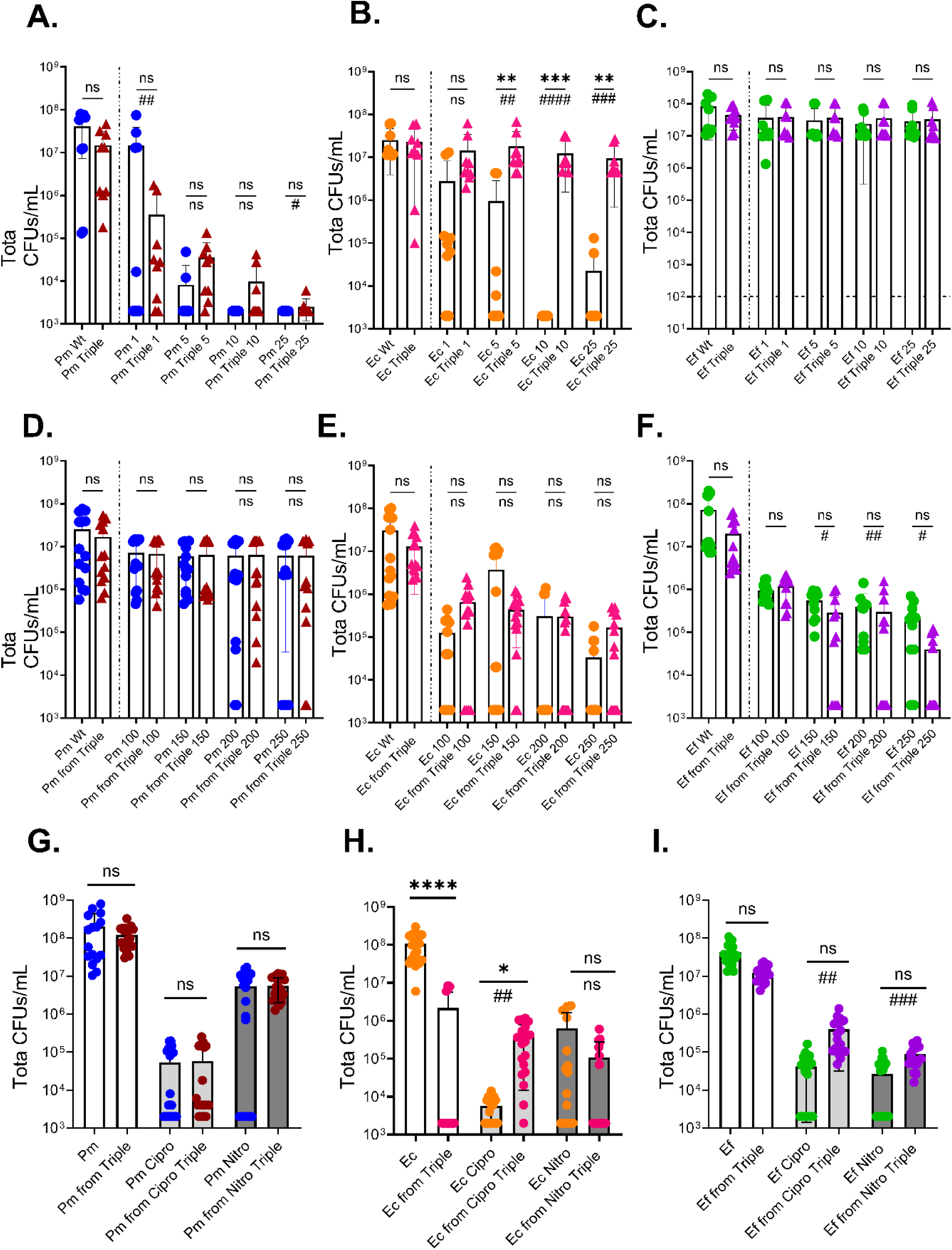
Polymicrobial biofilms provide species-specific protection from commonly used antibiotics. Single and triple species biofilms were established for 24 hours in diluted urine and exposed to (A-C) ciprofloxacin at 1, 5, 10, or 25 (D-F) nitrofurantoin at 100, 150, 200, 250 ug/mL. Data show 4 independent experiments with 2 biological replicates each. (G-I) Catheter biofilms were formed in the modified biofilm reactor in dilute urine for 24 hours and exposed to 25 ug/mL of ciprofloxacin or 200 ug/mL of nitrofurantoin. CFUs/mL were determined by differential plating. Data represent at least 3 independent experiments with at least 7 segments per catheter condition. Statistical significance was determined by Kruskal-Wallis test with multiple comparisons (asterisks) and by Chi-square analysis for number of catheter segments with CFUs above the limit of detection (pound signs); */#, P<0.05; **/##, <0.01; ***/###, P<0.001; ****/####, P<0.0001; ns, non-significant.

*E. coli* single species biofilms were sensitive to both ciprofloxacin and nitrofurantoin, and triple species biofilm formation provided robust and statistically significant protection from ciprofloxacin concentrations >5 ug/mL but no protection from nitrofurantoin (Figure 6B and E). *E. faecalis* single and triple species biofilms were completely resistant to ciprofloxacin (Figure 6C), and nitrofurantoin only resulted in a modest decrease in CFUs recovered from single species biofilms (Figure 6E). However, *E. faecalis* exhibited an unexpected further decrease in viability when triple species biofilms were treated with ≥150 ug/ml nitrofurantoin, as a greater percentage of catheter segments from the triple species biofilms exhibited CFU counts at or below the limit of detection (Figure 6F). These data indicate that while polymicrobial biofilm formation may provide protection to some species, this additional stressor may also influence polymicrobial interactions in a way that could increase susceptibility of others.

To determine whether flow conditions influence the antimicrobial susceptibility of each species in polymicrobial biofilms, the modified flow reactor was used to form single and triple species catheter biofilms in dilute human urine for 24 hours, followed by continued exposure to either untreated urine, urine with 25 ug/mL of ciprofloxacin, or urine with 200 ug/mL of nitrofurantoin for 24 hours. Biofilms CFUs were then derived from multiple 20 mm segments across the length of the catheter. *P. mirabilis* viability was more substantially reduced by ciprofloxacin than nitrofurantoin, and no significant differences were observed when comparing CFUs from the triple species biofilms and the single species biofilms formed (Figure 6G). However, 100% of the triple species catheter segments from the nitrofurantoin treatment had high *P. mirabilis* CFUs compared to only 57% of the single-species catheters, suggesting that triple species biofilm formation may provide some degree of nitrofurantoin protection to *P. mirabilis* under flow conditions. *E. coli* from untreated conditions had a decrease in CFUs while in triple species catheters (Figure 6H), which is again likely due to the prolonged impact of *P. mirabilis* urease activity at the 48 h time point. For ciprofloxacin treatment, triple species biofilm formation provided substantial protection to *E. coli,* resulting in higher CFUs compared to single species biofilms*. E coli* biofilms formed under flow were relatively susceptible to nitrofurantoin, although triple species biofilm formation did not provide any protection against this antibiotic. *E. faecalis* from untreated single and triple species both had a high level of CFUs recovered from the segments (Figure 6I). While triple species biofilm formation did not alter the total viability of *E. faecalis,* it did impact the likelihood of having high survival of *E. faecalis* within the biofilm: after treatment with either antibiotic: high CFUs of *E. faecalis* were only present on 66% and 52% of single species catheter segments after treatment, respectively, while 100% of segments from polymicrobial biofilms had high *E. faecalis* CFUs after treatment (Figure 6I).

### Triple species biofilm enhancement is driven by an increase in total protein

To further examine the underlying source of the increased biofilm biomass caused by co-culture of uropathogens, biofilm composition was characterized by assessing three common components of a bacterial biofilm: total protein content, extracellular DNA (eDNA), and carbohydrates. The abundance of eDNA, carbohydrates and protein was first measured from 6 pooled replicate biofilms formed in dilute human urine in 24 well plates to determine whether enhanced biofilm biomass corresponds to any of these components. No differences were observed in eDNA levels between the biofilms, with the exception that *E. faecalis* single species biofilms exhibited lower levels of eDNA than the other species (Figure 7A). Carbohydrate quantities were very low and therefore generally too close to the limit of detection for high confidence, but no differences were observed between single and polymicrobial biofilms (Figure 7B). However, total protein levels were significantly higher in all polymicrobial biofilms (Figure 7C). Biofilms formed in AUM and human urine exhibited similar trends (Supplemental Figure 3), although high background levels and variability were noted for protein quantification in human urine that abrogated statistical significance. Taken together, these data suggest that the increase in biofilm biomass for double and triple species biofilms is due at least in part to an increase in protein content in AUM and dilute human urine but may also be influenced by other as-yet unidentified factors in undiluted human urine.

**Figure 7:**
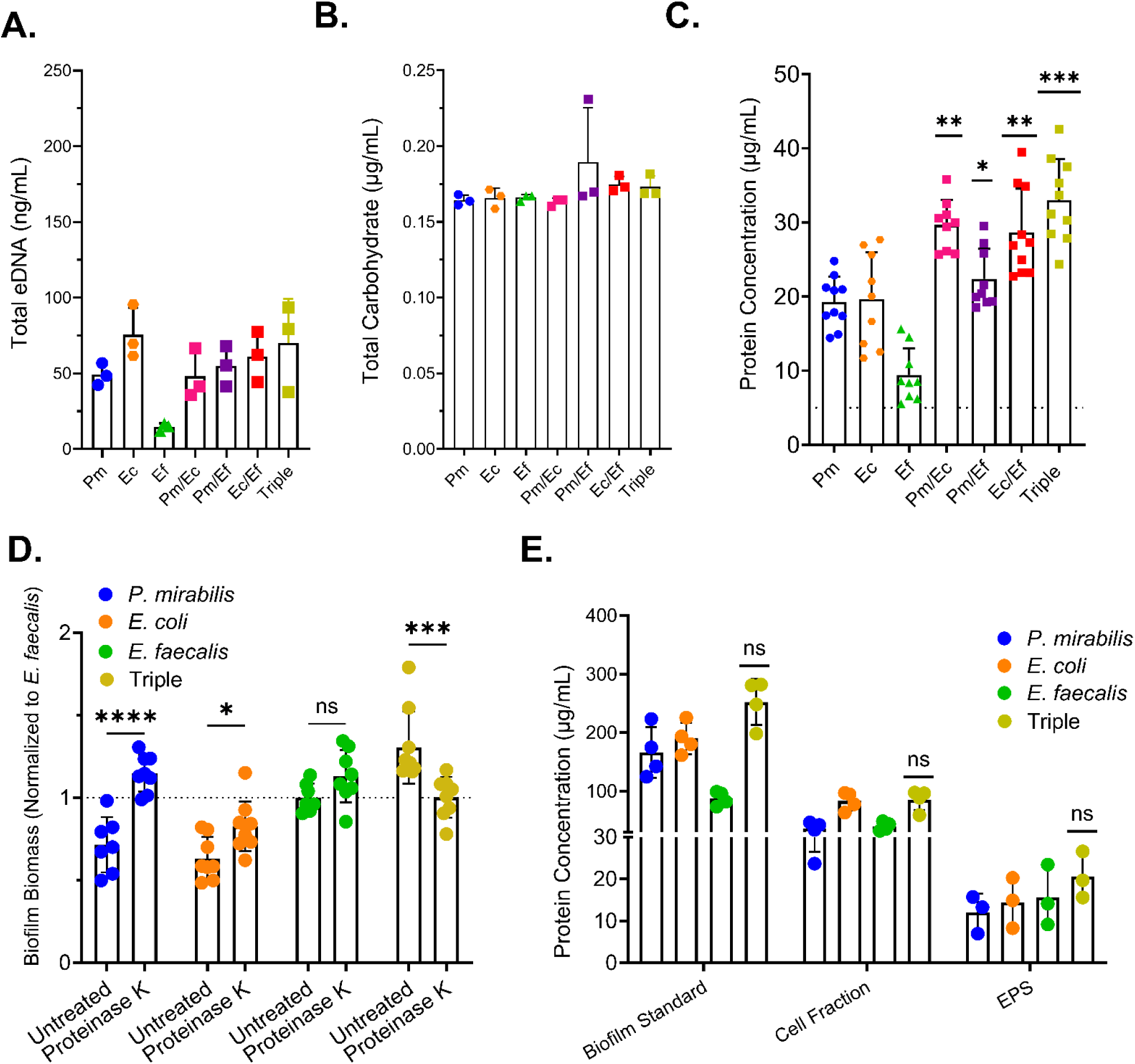
Triple species biofilm biomass enhancement is driven by an increase in total protein. Single, double, and triple species biofilms were established in 24 well plates for 24 hours in diluted urine. (A) Total eDNA concentration in single, double, and triple species biofilms. (B) Total carbohydrate concentration in single, double, and triple species biofilms. (C) Total protein concentration in single, double, and triple species. Polymicrobial combinations were compared to the highest associated single species concentration. Error bars represent means and SD from at least 3 biological replicates with at least 3 replicates each. Statistical significance was determined by an ordinary one-way ANOVA test with multiple comparisons; *, P<0.05; **, <0.01; ***, P<0.001; ns, non-significant. (D) Single and triple species biofilms were formed in diluted urine with or without proteinase K (30ug/mL) for 24 hours and then stained with crystal violet. The data represent three independent experiments with at least two biological replicates each. Statistical significance determined by 2way ANOVA with multiple comparisons test;*, P<0.05; ***, P<0.001; ; ****, P<0.0001 ns, non-significant. (E) Biofilms were separated into a cell-associated fraction and the EPS fraction, and protein concentration was analyzed by BCA from three independent experiments for single and triple species biofilms.

To further confirm that enhancement is driven by increased protein levels within the biofilm, single and triple species biofilms were established in diluted urine with and without proteinase K. The addition of proteinase K appeared to slightly increase biomass of the single species biofilms, as previously observed for *E. faecalis* (26), potentially due to the enzyme being incorporated into the biofilm or degraded for use as a nutrient. However, the presence of proteinase K abrogated the enhancement of the triple species biofilm (Figure 7D). These data further support the hypothesis that biomass enhancement is likely due to an increase in protein content.

To determine if the enhancement-associated increase in total protein derives from biofilm-associated bacteria or from the extracellular polymeric substance (EPS), biofilm fractionation was performed on single and triple species biofilms established in diluted urine. The eDNA, carbohydrate, and protein content of the unfractionated biofilm, the cell-associated fraction, and the soluble EPS fraction were then examined as above. As previously observed, no differences in carbohydrate or eDNA concentrations were seen between single and triple species biofilms in any fraction (Supplemental Figure 4). However, protein concentrations in the fractionated single and triple species samples demonstrated a trend towards being increased in the EPS rather than the cell-associated fraction (Figure 6C).

### L-fucose metabolism by *E. coli* supports triple species biofilm enhancement

Cooperation between *P. mirabilis*, *E. faecalis*, and *E. coli* have been previously investigated and demonstrated to occur in double species conditions. For example, ornithine export from *E. faecalis* via the ArcD antiporter was found to support arginine biosynthesis in *P. mirabilis* via ArgF, and this cross feeding mechanism contributed to increased biofilm biomass and protein content as well as infection severity in a mouse CAUTI model (11, 26). Ornithine export by *E. faecalis* ArcD was also shown to mediate beneficial interactions with *E. coli* by increasing siderophore production (enterobactin, via EntB) and ultimately increasing biofilm formation and infection under iron-limited conditions (23). We therefore investigated the contribution of *E. faecalis* ornithine export, *P. mirabilis* arginine biosynthesis, and *E. coli* enterobactin production to triple species biofilm enhancement. Surprisingly, triple species biofilms formed by any combination of mutants disrupted for each of these pathways still exhibited enhanced biomass (Supplemental Figure 5). These data suggest that triple species biofilm enhancement occurs independently of these known interactions.

To identify specific proteins that are enriched under conditions where polymicrobial biofilms have enhanced biomass, proteomics analysis was performed on single, dual, and triple species biofilms that were established in diluted urine in 24 well plates. Only proteins that could be uniquely associated with an individual species were included in the final analysis of relative abundance, in order to have sufficient confidence in the source of enrichment. For each species, protein abundance in dual and triple species biofilms were compared to the relevant single species biofilm to identify differences in relative abundance across conditions.

The protein profiles of each species were substantially altered in polymicrobial biofilms (Supplemental Figure 6). Since priority effect studies identified *E. coli* as the key mediator of triple species biofilm enhancement, we sought to focus on *E. coli* proteins that were specifically enriched under the triple species biofilm condition but not in either dual species biofilm. A total of 2450 proteins could be specifically mapped to *E. coli,* of which 194 (7.9%) exhibited a statistically-significant increase in abundance during polymicrobial biofilm formation (Figure 8A): 100 proteins (51%) were enriched during dual species biofilm formation with *E. faecalis,* 62 (32%) during biofilm formation with *P. mirabilis,* and 95 (49%) during triple species biofilm formation, with only 5 proteins (2.6%) enriched under all three conditions.

**Figure 8:**
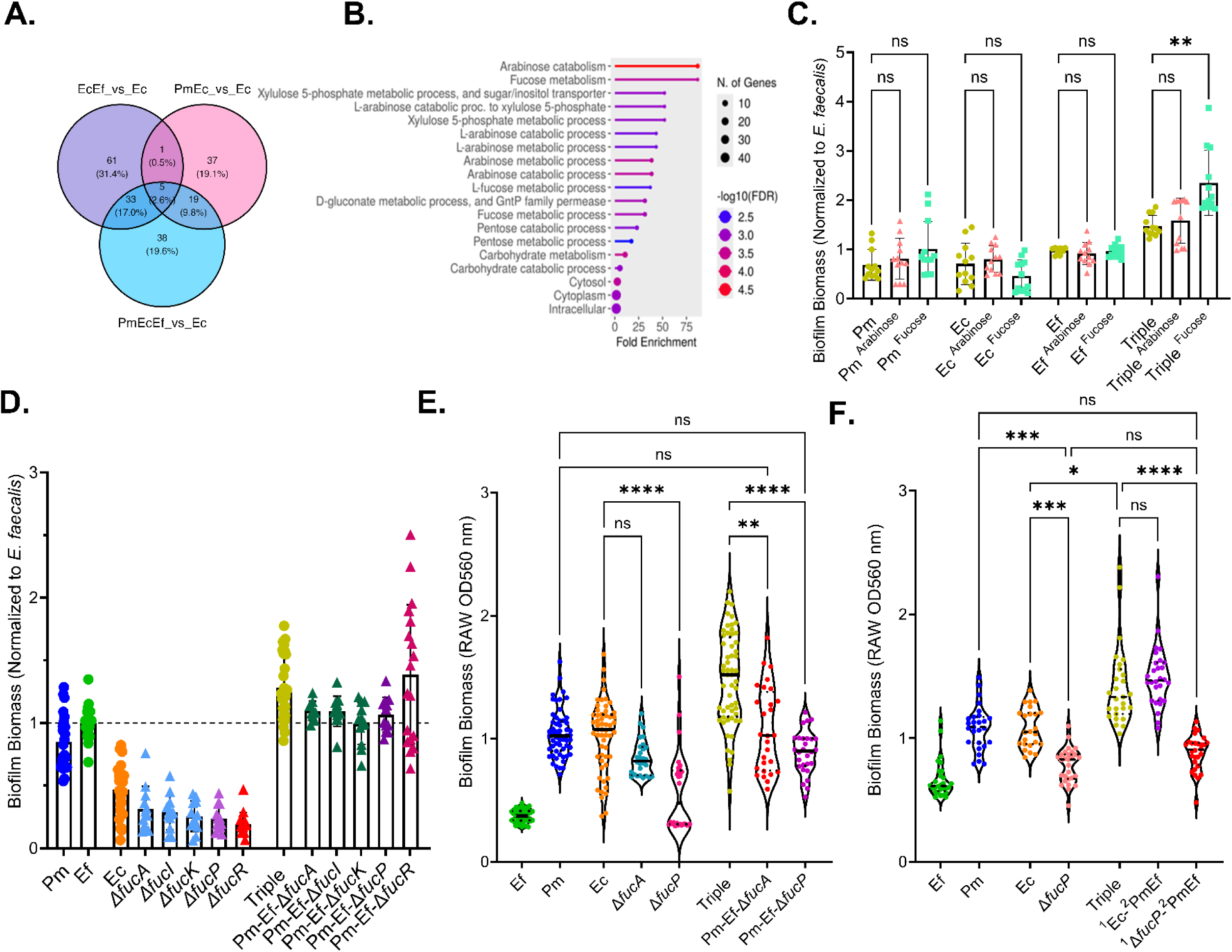
L-Fucose utilization by *E. coli* contributes to triple species biofilm enhancement. (A) Venn diagram demonstrating the overlap in *E. coli* proteins that were enriched in dual and triple species biofilms. (B) Pathway analysis of enhanced *E. coli* proteins that were specifically enriched in triple species biofilms. (C) Biofilm biomass quantification (crystal violet staining) of single and triple species biofilms established in 24 well plates in dilute human urine and supplemented with 10 mM L-arabinose or L-Fucose. Data show mean and standard deviation for 4 independent experiments with 3 biological replicates each. Statistical significance determined by one-way ANOVA with multiple comparisons; **, <0.01; ns, non-significant. (D) Single and triple species biofilms with mutants in *E. coli* related to L-fucose utilization. (E) Triple species biofilms under flow with *fucA* (red) and *fucP* (purple) mutants. Data shows 2 independent experiments with 14 segments per catheter condition and represent raw OD values. (F) 48 hour colonization priority experiments under flow conditions comparing the ability of *fucP* and wild-type *E. coli* to contribute to triple species biofilm enhancement. Data represent 2 independent experiments with14 catheter segments per condition. Statistical significance determined by Kruskal-Wallis with multiple comparisons; *, P<0.05; ***, P<0.001; ****, P<0.0001; ns, non-significant.

Pathway analysis of the *E. coli* proteins that were specifically enriched in triple species biofilms revealed numerous proteins involved in arabinose, fucose, and xylulose metabolic processes (Figure 8B). To determine if either L-arabinose or L-fucose can directly alter biofilm biomass, single and triple species biofilms were formed in diluted urine with or without 10 mM arabinose or fucose for 24 hours and biomass was examined by crystal violet staining. L-arabinose had no impact on biofilm biomass, but L-fucose supplementation significantly enhanced triple species biofilm biomass without impacting any of the single species biofilms (Figure 8C). Interestingly, this enhancement was specific to the triple species condition as fucose supplementation had no impact on dual species biofilms (Supplemental Figure 7). Thus, fucose appears to play an important role in mediating biofilm-enhancing polymicrobial interactions only in the presence of all three species.

Fucose utilization in *E. coli* is performed by the *fucPIK* operon (transport and isomerization) and the adjacent *fucAO* operon (catabolism), and expression of both operons is tightly regulated by the global regulator Crp as well as FucR (41, 42). To determine whether fucose utilization is necessary for triple species biofilm enhancement, mutants in *fucA,fucI,fucK,fucP,* and *fucR* were used. None of the fucose mutants exhibited significant alterations in single species biofilm biomass compared to wild-type *E. coli* (Figure 8D). However, when used to establish triple species biofilms, enhancement was lost for all except *fucR,* confirming that fucose utilization by *E. coli* is necessary for triple species biofilm enhancement (Figure 8D).

To determine whether fucose utilization also contributes to catheter biofilm enhancement under flow conditions, single and triple species biofilms were established in the modified biofilm flow reactor in diluted urine for 24 hours (Figure 8E). Single species biofilms formed by Δ*fucA* and Δ*fucP* exhibited a reduction in biomass as compared to wild type *E. coli*, stressing the importance of L-fucose utilization for *E. coli* biofilm formation under flow conditions. Triple species biofilms formed with either Δ*fucA* or Δ*fucP* also exhibited a clear loss of biofilm enhancement, as compared to the *P. mirabilis* single species biofilms as well as the triple species biofilm formed with wild type *E. coli* (Figure 8E).

We next examined the contribution of L-fucose utilization to the ability of *E. coli* to create a permissive environment for polymicrobial biofilm enhancement by establishing single species biofilms with either wild-type *E. coli* or the *fucP* mutant in the flow model for 24 hours and then co-inoculating with *P. mirabilis* and *E. faecalis* (Figure 8F). Single species biofilms formed by the *fucP* mutant had significantly lower 48 hour biomass than those formed by wild-type *E. coli,* but comparable to that of *E. faecalis*. However, loss of *fucP* completely abrogated the ability of *E. coli* to support triple species biofilm enhancement, as the triple biofilm formed by initial seeding with *fucP* remained at a comparable level of biomass as the *fucP* mutant single species biofilm (Figure 8F). Thus, fucose utilization by *E. coli* contributes to the establishment of a permissive environment on the catheter surface that facilitates polymicrobial biofilm formation by *P. mirabilis* and *E. faecalis* and ultimately enhances biofilm biomass over what can be achieved by any dual-species combination.

## Discussion

Asymptomatic colonization of urinary catheters frequently occurs during long term use of these medical devices, and the resulting bacterial communities can become highly persistent and recalcitrant to antibiotics. Recent studies have found that *P. mirabilis*, *E. coli*, and *E. faecalis* are the most common and persistent co-colonizing partners of the catheterized urinary tract, suggesting potential synergistic interactions. This study aimed to examine the relationship between *P. mirabilis*, *E. coli*, and *E. faecalis,* specifically focusing on how polymicrobial interactions influence the development of antibiotic recalcitrant catheter biofilms. Overall, our findings suggest that these three uropathogens exhibit a synergistic partnership that enhances biofilm biomass, and that L-fucose metabolism by *E. coli* is critical for creating the initial environment that facilitates this partnership.

Free L-fucose in urine exists in diminutive amounts of ∼16 ug/mL (97.5 uM) in healthy individuals, and fucosuria (high levels of fucose in urine) has been linked to markers of disease such as bladder cancer and liver disease (43–45). However, the amount present in our pooled diluted urine is clearly sufficient for metabolism by *E. coli,* since disruption of fucose utilization resulted in a biofilm defect in the catheter flow model and prevented *E. coli* from establishing a permissive environment for triple species biofilm enhancement when inoculation order was staggered. The downstream effects of fucose metabolism on *E. coli* biofilm formation and composition still remain to be determined. One possibility is that fucose metabolism may allow

*E. coli* to alter the nutrient composition of the urine in a way that supports adherence and biofilm production by *P. mirabilis* and *E. faecalis.* Alternatively, fucose metabolism may impact production of *E. coli* adhesions or EPS components that then facilitate adherence and biofilm formation by *P. mirabilis* and *E. faecalis*.

To our knowledge, no prior studies have reported a role for L-fucose in mediating biofilm formation by uropathogens or for mediating polymicrobial interactions. However, others have demonstrated species-specific effects of L-fucose metabolism on monomicrobial colonization and biofilm formation in infection niches. In *Klebsiella pneumoniae*, the metabolism of L-fucose was found to promote gastrointestinal colonization and production of virulence factors (46). In minimal media, the addition of 0.5% L-fucose was sufficient to increase biofilm biomass and impacted the production of type III fimbriae. Similar to our study, L-fucose increased total protein content in the *K. pneumoniae* biofilms without altering other biofilm components. The contribution of L-fucose to biofilm formation and architecture may therefore be similar between

*E. coli* and *K. pneumoniae.* Further study is needed to determine the impact of L-fucose on production of adhesins in both species, and to determine if *K. pneumoniae* may similarly facilitate polymicrobial biofilm formation. In contrast, fucose utilization decreased biofilm formation and enhanced chemotaxis in *Campylobacter jejuni* (47). When *C. jejuni* was cultured in Mueller Hinton Broth (MH) supplemented with 25 mM fucose, a reduction in biofilm biomass was observed as compared to the control group lacking supplementation. Interestingly, the loss of *ΔfucP* in *C. jejuni* did not impact biofilm formation in the presence or absence of fucose supplementation, suggesting that L-fucose utilization is dispensable for this bacterium and that accumulation may impair biofilm formation.

Once within a polymicrobial setting, *E. coli* may also be able to scavenge fucose from other bacterial species. While some genomes of *Proteus mirabilis* harbor fucose synthase (*fcl*) for conversion of mannose to fucose, the closest match found by BLAST in *P. mirabilis* strain HI4320 was only 24% amino acid identity. Similarly, several *Enterococcus faecalis* genomes harbor *fcl* but BLAST did not reveal any homologs in strain OG1RF. Thus, it is unclear whether either of the isolates used in our study may be capable of producing L-fucose to further fuel metabolism in *E. coli.* However, since biofilm enhancement in combinations involving *E. coli* was only consistently observed in the triple species condition and not in any of the dual species biofilms, our data suggest that enhancement ultimately requires a specific interaction between both *P. mirabilis* and *E. faecalis* with *E. coli* that would not be driven by fucose production from one species alone. The only exception was that dual species biofilms of *E. coli* with *E. faecalis* exhibited enhancement in AUM, suggesting that differences in growth media influence metabolic interplay between these species to support polymicrobial biofilm formation. The observation that ornithine/arginine-mediated interactions between *P. mirabilis* and *E. faecalis* that are sufficient to drive dual-species enhancement are no longer required when *E. coli* is present provides further support for multiple condition-specific mechanisms underlying biofilm enhancement.

L-fucose in the intestinal tract is known to regulate virulence for pathogens like Enterohaemorrhagic *E. coli* (EHEC), *Salmonella typhimirium,* and *Clostridium difficile,* but this requires the cooperation of members of the gut microbiota like *Bacteroides thetaiotaomicron* to release fucose from epithelial cells (48–53). In EHEC, the environmentally free L-fucose is sensed by a histidine kinase response regulator, FusKR, which then represses virulence related factors and provides a growth advantage in the lumen in a mouse model (49, 54). Host adrenergic signals and microbiota-generated autoinducer signals were found to repress expression of FusKR, and subsequently enhanced EHEC virulence (55, 56). The mucosal surface of the bladder may similarly provide an environment form which fucose could be scavenged, and this could potentially be mediated by members of the urinary microbiome in a similar manner as members of the but microbiota (57–61). However, little is known regarding the interaction of the urinary microbiota with the bladder mucosal surface. Considering that our proteomics results did not show an enrichment of *E. coli* fucosidases or histidine kinases under any condition and there is no mucosal surface under our experimental conditions, it is unlikely L-fucose is driving triple species biofilm formation in a similar manner as what has been described for EHEC. What does remain evident is that L-fucose utilization by *E. coli* supports colonization and could be supporting polymicrobial relationships, even on an abiotic surface in the absence of host cells. Thus, further exploration is warranted to dissect the exact mechanism by which this metabolite facilitates polymicrobial biofilm formation.

Catheter-associated bacteriuria (CA-ASB) is largely polymicrobial and asymptomatic for the host and can persist for long periods of time. Thus, examining mechanisms that contribute to polymicrobial biofilm formation may provide insight into how CA-ASB communities form and remain asymptomatic, as well as novel points of intervention to disrupt polymicrobial biofilm formation and improve efficacy of antibiotic treatment when needed. Biofilm communities that form on the catheter may function as a unique biome that largely favors stable colonization rather than the transition to invasive disease. Observations from the catheter flow model support this idea, since initial colonization by *E. coli* facilitated formation of the robust triple species biofilm without outgrowth by a dominate species in polymicrobial biofilms. Enhancement of triple species biofilms form with patient isolates likewise suggests that this interaction does occur within the catheterized host and across multiple different clinical isolates.

However, when conditions in the urinary tract are disturbed, the individual members of this biome may shift to a pathogenic lifestyle, promoting tissue invasion, ascension to the kidneys, and severe disease. Disruption of the urinary biome, the damage to the mucosal layer, and damage to bladder epithelial cells by long term catheter usage could therefore trigger outgrowth or a virulent phenotype in pathogenic members of the microbial community. Further research is therefore needed to fully dissect the interactions between uropathogens within polymicrobial catheter biofilms and determine whether L-fucose mediated interactions could represent a target either for disrupting persistence or facilitating re-establishment of asymptomatic colonization after a perturbation rather than the transition to invasive disease.

## Methods

### Bacterial strains

*Proteus mirabilis* HI4320, *Providencia stuartii* BE2467, and *Morganella morganii* TA43 are well-established and minimally-passaged clinical isolates from the urine of long-term catheterized patients in a chronic care facility (62). *Enterococcus faecalis* OG1RF is a well-established oral clinical isolate commonly used in CAUTI research (63, 64)*. Escherichia coli* CFT073 was isolated from a patient with chronic acute pyelonephritis and is also commonly used in UTI and CAUTI research (65). *P. mirabilis ureF* refers to a mutant of *P. mirabilis* HI4320 in which the gene for the UreF accessory protein of the urease operon was disrupted by insertion of a kanamycin cassette using the TargeTron protocol, resulting in loss of urease activity (66). *E. coli* CFT073 fucose utilization mutants *fucA,fucI,fucK,* and *fucP* were obtained from an ordered library of transposon mutants (67). Staphylococcus aureus UTI MRSA was isolated from chronically infected individuals with indwelling urinary catheters (6, 68).

### Bacterial culture conditions

*P. mirabilis* and *E. coli* were cultured in 5 mL of low-salt Luria-Bertani (LSLB) broth (10 g/L tryptone, 5g/L yeast extract, 0.1 g/L NaCl) at 37°C with shaking at 225 RPM, while *E. faecalis* was cultured overnight in Brain Heart Infusion (BHI) broth at 37°C with shaking at 225 RPM. Bacteria were also cultured in other media types as indicated, including tryptic soy broth supplemented with 1% glucose (TSBG) and modified artificial urine medium (AUM) following the Brooks recipe (69). Filter-sterilized pooled human urine was purchased from Cone Bioproducts (Sequin, TX), stored at −20°C, and in some assays was diluted 1:1 with 0.9% saline to adjust the specific gravity to 1.004. Where indicated, dilute urine was supplemented with 30 ug/mL proteinase K (MP Biomecials,C183988, LU1124039730-1), 10 mM arabinose (MP Biomedicals,C100706, LU1124156894-1), or 10 mM fucose (Thermo, CA16789.03, LR0BJ007). For differential plating, *P. mirabilis* was incubated on either LSLB agar supplemented with 2.5 ug/mL tetracycline or MacConkey agar, *E. faecalis* was grown on BHI agar supplemented with 100 ug/mL of streptomycin, and *E. coli* was plated on MacConkey agar for enumeration by colony morphology or on plain LSLB, with enumeration by subtracting the CFUs of *P. mirabilis* on LSLB+tet from the total CFU/ml of both *P. mirabilis* and *E. coli* on LSLB..

### Planktonic growth curves

Overnight cultures of all species were adjusted to ∼10^6^ CFU/mL in the indicated media (TSBG, AUM, urine, diluted urine). Single species cultures were seeded with 1×10^6^ CFU/ml of the indicated species, dual species cultures were seeded with ∼5 x10^5^ CFU/ml of each respective species, and triple species cultured were seeded with ∼3.3 x 10^5^ CFU/ml of each species, such that the total number of viable bacteria for all cultures was 1×10^6^ CFU/ml. 5mL cultures were incubated at 37°C with shaking at 225 rpm, and aliquots were taken hourly from 0-8 hours as well as 24 hours, serial diluted, and plated as described above and using an EddyJet 2 spiral plater (Neutec Group) for the determination of CFUs using a ProtoCOL 3 automated colony counter (Synbiosis).

**Crystal violet staining of bacterial biofilms.**

Overnight cultures of bacteria were adjusted to ∼10^6^ CFU/mL as described above in the indicated media and 750 uL was dispensed into triplicate wells in tissue culture treated 24-well plates (Falcon 353047). Three additional wells were filled with sterile media to serve as blanks for crystal violet staining. Plates were incubated for 24 hours at 37°C in a partially sealed plastic bag with a damp paper towel to maintain humidity. After 24 hours, supernatants were gently aspirated and wells were washed twice with 1mL of 1x phosphate buffer saline (PBS). Biofilms were fixed by adding 1 mL of ice cold 95% ethanol to each well and incubating for 15 minutes at room temperature (RT).

Ethanol was then aspirated and the wells were dried for 30 minutes in a fume hood. A 0.1% crystal violet solution (Fisher, CS252275A) was added to the dried wells and incubated at RT for 60 minutes to stain all biomass. Wells were then aspirated, washed with 1 mL of deionized water, and solubilized in 1 mL of 95% ethanol with shaking at 220 RPM for 15 minutes. Wells were then scraped with a 1000 uL pipette tip to ensure full resuspension of all biomass in EtOH.Crystal violet absorbance read at 570 nm using a BioTek synergy H1 plate reader in a a 96-well plate, and samples were typically diluted 1:5 in EtOH to ensure an accurate reading. For most experiments, crystal violet absorbance is expressed relative to single species *E. faecalis* biofilm unless stated otherwise to facilitate comparison across independent experiments.

### Determination of cell viability within biofilms

Biofilms were established in triplicate in tissue culture treated 24-well plates as described above and incubated for 24 hours at 37°C. Wells were washed with 1 mL of sterile 1X PBS and were scraped as described above for resuspension using a sterile pipette tip. Samples were serially diluted and plated on appropriate agar for differentiation of each species as described above using an EddyJet2 spiral plater (Neutec Group) to determine CFUs using a ProtoCOL 3 automated colony counter (Synbiosis).

### Protein, eDNA, and carbohydrate analyses

24 well plates were inoculated and incubated as described above, with 6 replicate wells per inoculum. Supernatants were carefully aspirated and biofilms were gently washed with 1X PBS. The first well for a given inoculum was then resuspended in 1X PBS and scraped as described above to collect all biofilm biomass. This resuspension was then transferred to the next well and repeated until all 6 replicate wells had been resuspended together in 1ml of PBS. The protein concentration of each biofilm suspension was measured using a Bicinchoninic acid assay (BCA) assay (Thermo, P23252) following the manufacturer’s instructions A Quant-iT dsDNA high sensitivity kit (Thermo, P11496) was used to measure double-strained DNA, and carbohydrate measurements were performed using a kit (Sigma Aldrich, MAK559) with a D-glucose ladder.

### Silicone disc biofilms

Silicone disc were punched from a sheet using a 9 mm dermal punch and sterilized before use by autoclaving. Discs were then aseptically deposited into the wells of a non-tissue culture treated 24 well plate (Falcon, 351147), and biofilms were established by inoculating as described above After 24 hours, silicone discs were carefully removed with forceps, gently washed with 1X PBS, and air dried overnight. Discs were then stained with 1 mL of 0.1% crystal violet solution in a 1.5 mL Eppendorf tube for 15 minutes, washed in ddH2O twice to remove excess stain, and placed into 1mL of 95% ethanol. Samples were vortexed for 15 minutes, 25 µL of each ethanol suspension was transferred to a 96-well and diluted 1:5 with EtOH, and absorbance was read at 570 nm as described above. For assessment of viable bacteria within the silicone disc biofilms, the silicone disc biofilms were removed from the well, washed with 1X PBS, and transferred to Eppendorf tubes containing 1 mL of 1X PBS, vortexed for 15 minutes, serially diluted, and plated as described above.

### Glass bladder biofilms

A previously described glass bladder flow model was used, in which a 500 mL water-jacketed vessel is maintained at 37°C by a circulating water bath. A Foley catheter is inserted, inflated, and urine is supplied into the bladder at a constant flow rate of 0.75-1.0mL/min through a peristaltic pump. CFUs and crystal violet analysis were performed as described previously.(68)

### Catheter biofilm reactor

The periodic flow catheter model was based off design instructions from the construction of a drip biofilm reactor (70).The biofilm reactor consists of two boxes; the outer box is Styrofoam with a heating mat (BN-LINK, 5021701) on the inside, which is kept at 37°C and monitored by a thermal probe. The inner box is polypropylene and used to hold the catheters in place and ensure temperature regulation. See Supplemental Figure 2 for diagram.

French 14 catheters (Dynarex, 5075/57758) are first exposed to undiluted pooled human urine to establish a sterile conditioning film to facilitate bacterial attachment. To do this, 3 mL of human urine is pushed through the lumen at the base catheter via 5 mL Luer-Lok tip syringe and a tubing clamp is placed underneath the eyelet to create a seal. The clamped catheter with the syringe attached is then incubated overnight at 37°C. The clamp is then removed and the attached syringe is used to aspirate the urine from the catheter lumen. Any residual urine is then removed by a vacuum pump. The top portion of the catheter (eyelet and balloon) are then removed using a sterile surgical scalpel to expose the lumen for inoculation.

Bacterial inocula were prepared to ∼10^6^ CFU/mL as described above in a final volume of 3 mL of dilute urine. Catheters are seeded at the base using a 5 mL syringe. After, a tubing clamp is placed at the base while a 14-gauge need is inserted into the end of the tubing where the balloon and eyelet were removed. The bacterial inocula are incubated for 1.5 hours at 37°C to promote bacterial attachment to the condition film within the catheter lumen. After incubation, catheters are fed through the biofilm reactor box and attached to a peristaltic pump (VWR/Masterflex) via the needle end of the catheter. After attachment, the clamp is removed and the drainage port end of the catheter is placed over a waste vessel. Urine flow is then established via peristaltic pump at 0.8 RMP for 1.5 minutes to clear the inocula with fresh diluted urine. Fresh dilute urine is then supplied every hour for 1.5 minutes (∼ 2mL of urine per cycle) for up to 24 or 48 hours, depending on the experiment.

For biofilm biomass quantification, catheters were gently aspirated by a vacuum pump to remove media from the lumen. Catheters were gently washed with 5 mL of ddH_2_O and dried in a biosafety cabinet for 24 hours. Catheter lumens were then stained with a 0.1% of crystal violet for 30 minutes, aspirated, and washed twice with 5 mL of ddH_2_O. The stained catheters were then sectioned into 20 mm segments using a scalpel and placed on a Kimwipe (Kimtech) to remove any excess liquid in the lumen. A single catheter provides 14 segments, which were each placed into individual 1.5 mL Eppendorf tubes with 1mL EtOH and vortexed for 15 minutes to fully solubilize. Crystal violet absorbance was then determined as previously stated. Raw OD values were reported rather than normalizing to *E. faecalis* due to the segment-to-segment variability in crystal violet staining. For CFU determination, catheters were washed as above, sectioned, and each section was vortexed in 1mL of 1X PBS for 15 minutes.

Suspensions were then diluted and plated as described above.

### Biofilm fractionation experiments

Single and triple species biofilms were established in 24 well plates as described above. The biomass from 6 wells per inoculum were scraped and pooled in 3 mL of Milli-Q H_2_O, and 1.5 mL of the biofilm sample was immediately frozen for use as unfractionated control while the remaining 1.5 ml was fixed with 21 µL of 16% formaldehyde and incubated with shaking at RT for 1 hour. For EPS extraction, 300 uL of 1M NaOH was added to samples and incubated at 37°C with shaking at 220 rpm for 90 minutes. To separate the soluble EPS from the cell-associated biofilm fraction, samples were centrifuged at 20,000xg for 1 hour at 4°C. Supernatants were transferred to a 2mL tube, while cell pellets were resuspended in 200 uL of Milli-Q H_2_O and frozen to represent the cell-associated fraction of the biofilm. The supernatants were further processed by passing through a 0.22 μm filter and removed of salts and other containments using dialysis cassettes (Thermo, P66110) in ddH_2_O overnight. Dialyzed samples were transferred to 2mL tubes and concentrated using a Speed Vac (Savant, VLPI 20) and reconstituted to a total volume of 200 uL with Milli-Q H_2_O and frozen to represent the EPS fraction of the biofilm. The biofilm sample, cell-associated fraction, and EPS portions of the samples were then subjected to BCA, carbohydrate, and eDNA analysis as described above.

### Antibiotic treatment

Single, polymicrobial biofilms were formed for 24 hours in diluted urine in the standard methods previously described. After incubation, using a vacuum pump supernatant was removed from biofilms, fresh diluted urine supplemented with 100, 150, 200, 250 ug/mL of nitrofurantoin (TCI, N0883, L:DBOTA-SH) or 1, 5, 10, 25,50 ug/mL of ciprofloxacin (C2510, L:SOBSA-BD) was then added to the wells. For biofilms underflow, antibiotic treatment was directly added to media bottle at 200 ug/mL for nitrofurantoin and 25 ug/mL for ciprofloxacin. Biofilms were exposed to the treated dilute urine for another 24 hrs; after biofilms were processed for CFUS using differential plating in the previously described manner.

### Proteomics

Sample preparation was from single, double, and triple species biofilms were prepared in 1 mL of ddH_2_O and from 6 pooled technical replicates. The mass spectrometry proteomics data have been deposited to the ProteomeXchange Consortium via the PRIDE partner repository with the dataset identifier PXD074670.

#### Protein Digestion

Fifteen micrograms of protein were taken from each sample, dried using a SpeedVac SPD 140DDA Vacuum Concentrator (Thermo Fisher, Scientific, San Jose, CA), and reconstituted in 30 µL of 50mM Tris-FA buffer (pH 8.5) containing 0.01% N-dodecyl-β-D-maltoside (DDM). Protein digestion was performed using a previously reported surfactant cocktail-aided extraction/precipitation/on-pellet digestion (SEPOD) protocol (71, 72). Briefly, proteins were first reduced with 5 mM dithiothreitol (DTT) at 56 °C for 30 min, followed by alkylation with 20 mM iodoacetamide (IAM) at 37 °C for 30 min in the dark. Both reduction and alkylation steps were carried out with constant mixing at 550 rpm using a Thermomixer. Protein precipitation was achieved by adding five volumes of chilled acetone, followed by vortexing, and incubation at −20 °C for 3 h. Samples were then centrifuged at 20,000 × g for 30 min at 4 °C, after which the supernatant was removed. The resulting protein pellets were washed with 200 µL methanol and air-dried. Pellets were subsequently reconstituted in 50 mM Tris-FA. Trypsin (Sigma-Aldrich), activated in 50 mM Tris-FA, was added, and enzymatic digestion was carried out at 37 °C for 16 h with constant mixing. Digestion was quenched by the addition of formic acid (FA) to a final concentration of 1%. Samples were centrifuged at 20,000 × g for 30 min at 4 °C. Two microliters from the supernatants of each sample were pooled to generate a quality control sample for assessing quantitative reproducibility, and the remaining supernatants were transferred to LC vials for LC-MS analysis.

Liquid chromatography-tandem mass spectrometry analysis:

Liquid chromatography-tandem mass spectrometry (LC-MS/MS) analysis was performed using a trapping nano-flow LC-Orbitrap Astral MS system, consisting of a Dionex Ultimate 3000 nano LC system, a Dionex Ultimate 3000 gradient micro-LC system equipped with a WPS-3000 autosampler, and an Orbitrap Astral mass spectrometer (Thermo Fisher Scientific, San Jose, CA).

For each sample, a single injection of 500 ng of digested peptides was analyzed. Three consecutive injections of pooled samples were also analyzed. A trapping column (150 µm i.d. × 5 cm, packed with 5 µm C18, CoAnn Technologies, Richland, WA) was used upstream of the nano-LC analytical column (75 µm i.d. × 20 cm, packed with 1.7 µm C18, CoAnn Technologies, Richland, WA) to remove matrix components and enable selective peptide delivery. Mobile phase A consisted of 0.1% formic acid (FA) in 2% acetonitrile, and mobile phase B consisted of 0.1% FA in 88% acetonitrile. The LC gradient was programmed as follows: 4-11%B for 2 min, 11-12%B for 2 min, 12-30% B over 26 min, 30-97% B over 0.1 min, followed by an isocratic hold at 97% B for 3 min. Mass spectrometric analysis was conducted in data-independent acquisition (DIA) mode. MS1 spectra were acquired over an m/z range of 380-980 at a resolution of 180,000, with a maximum injection time of 5 ms, and a normalized automatic gain control (AGC) target of 500%. Precursor ions were isolated using 2 Th-wide isolation windows and fragmented by higher-energy collisional dissociation (HCD) at a normalized collision energy of 25%. MS2 spectra were acquired over an m/z range of 350-1500 in the Astral (AST) mass analyzer, with a normalized AGC target of 200% and a maximum injection time of 3 ms.

#### Protein identification and quantification

The MS raw files were converted to HTRMS format using HTRMS Converter (Biognosys, Zurich, Switzerland) and searched against a combined NCBI reference sequence database for *Escherichia coli* CFT073, *Enterococcus faecalis* OG1RF, and *Proteus mirabilis* HI4320 using Spectronaut (version 19, Biognosys, Zurich, Switzerland) in DirectDIA mode. The protease was set to trypsin with up to two missed cleavages allowed. Carbamidomethylation of cysteine was specified as a fixed modification, while oxidation of methionine and protein N-terminal acetylation were specified as variable modifications. Protein-level false discovery rate (FDR) was controlled at 1% across the analysis. Only proteotypic peptides were used for protein quantification. All other parameters were kept at default settings.

#### Statistical analysis

Statistical analyses were performed using R (version 4.4.1). Species-specific normalization was carried out using median normalization. Comparisons betweengroups were conducted using two-sample, independent Student’s *t*-tests. Differentially expressed proteins were defined as those exhibiting a fold change greater than 1.3 or less than 0.77 with a p-value < 0.05.

#### Pathway Analysis

Differential protein abundance was calculated as log₂ fold change (Log₂ FC) comparing the triple-species biofilm to single biofilms. Proteins were considered significantly upregulated if they met both criteria: Log₂ FC > 1 (>2-fold increase) and p-value < 0.05. Proteins were identified using the NCBI RefSeq. Gene names were based on the STRING database and eggnog-mapper functional annotation tool. Pathway level analysis was done using the KEGG Ontology (KO) terms. Volcano plots were generated using Python (matplotlib) with log₂ fold change on the x-axis and −log₁₀(p-value) on the y-axis with threshold at Log₂ FC = 1 and p-value = 0.05.

### Statistical Analyses

Statistical significance from experimental results for biofilms, CFUs, and growth curves was assessed by one-way ANOVA with multiple comparisons or by two-way ANOVA as indicated by legends. Chi square tests were performed where indicated. These analyses were performed using GraphPad Prism, version 10.5.0.

## Acknowledgements

This work was supported by the National Institute of Diabetes and Digestive and Kidney Diseases under award R01 DK140371 to CEA and by the National Institute of Allergy and Infectious Diseases under award F31 AI183838 to SMT.

---

**Table 1:**
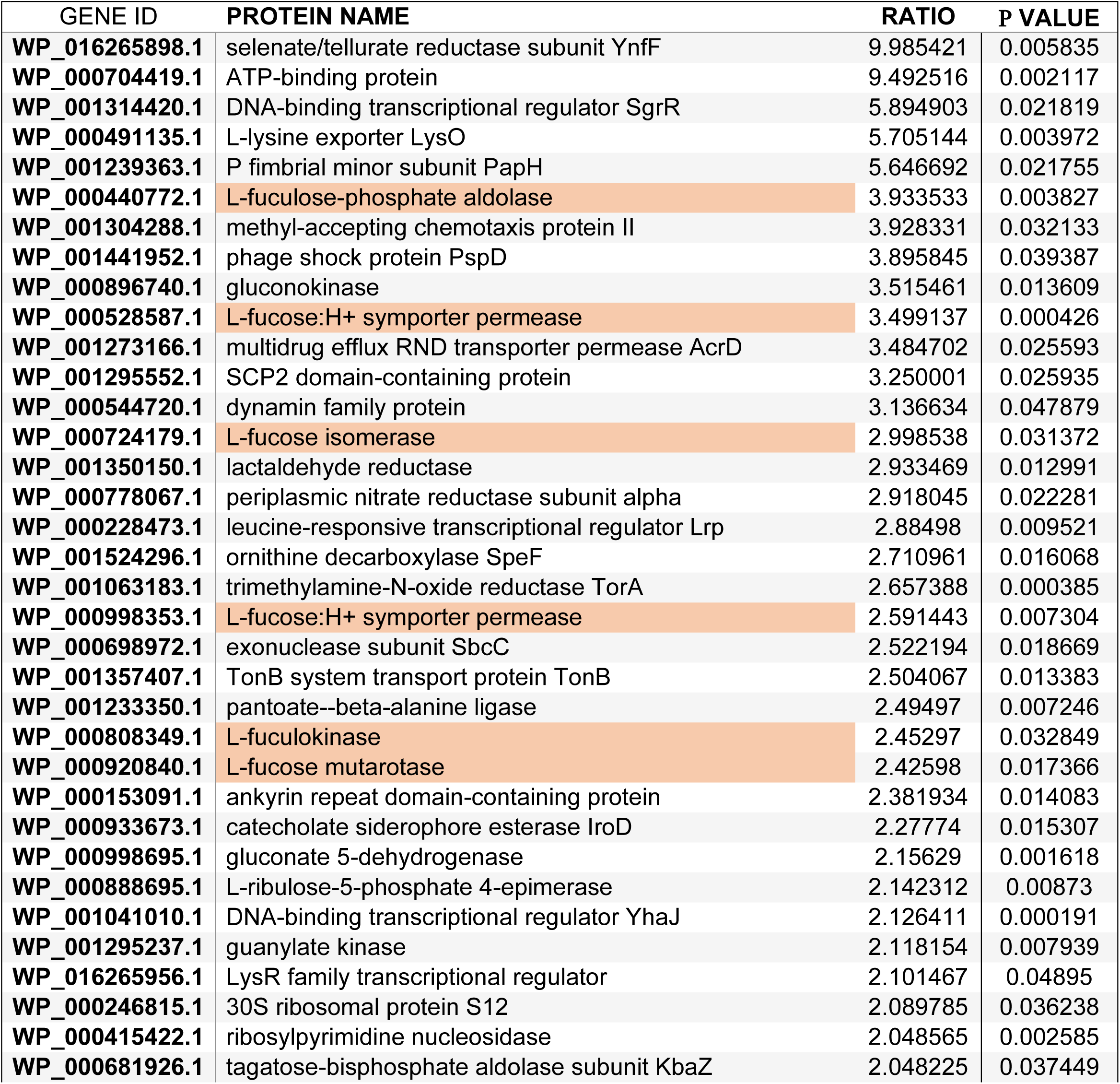
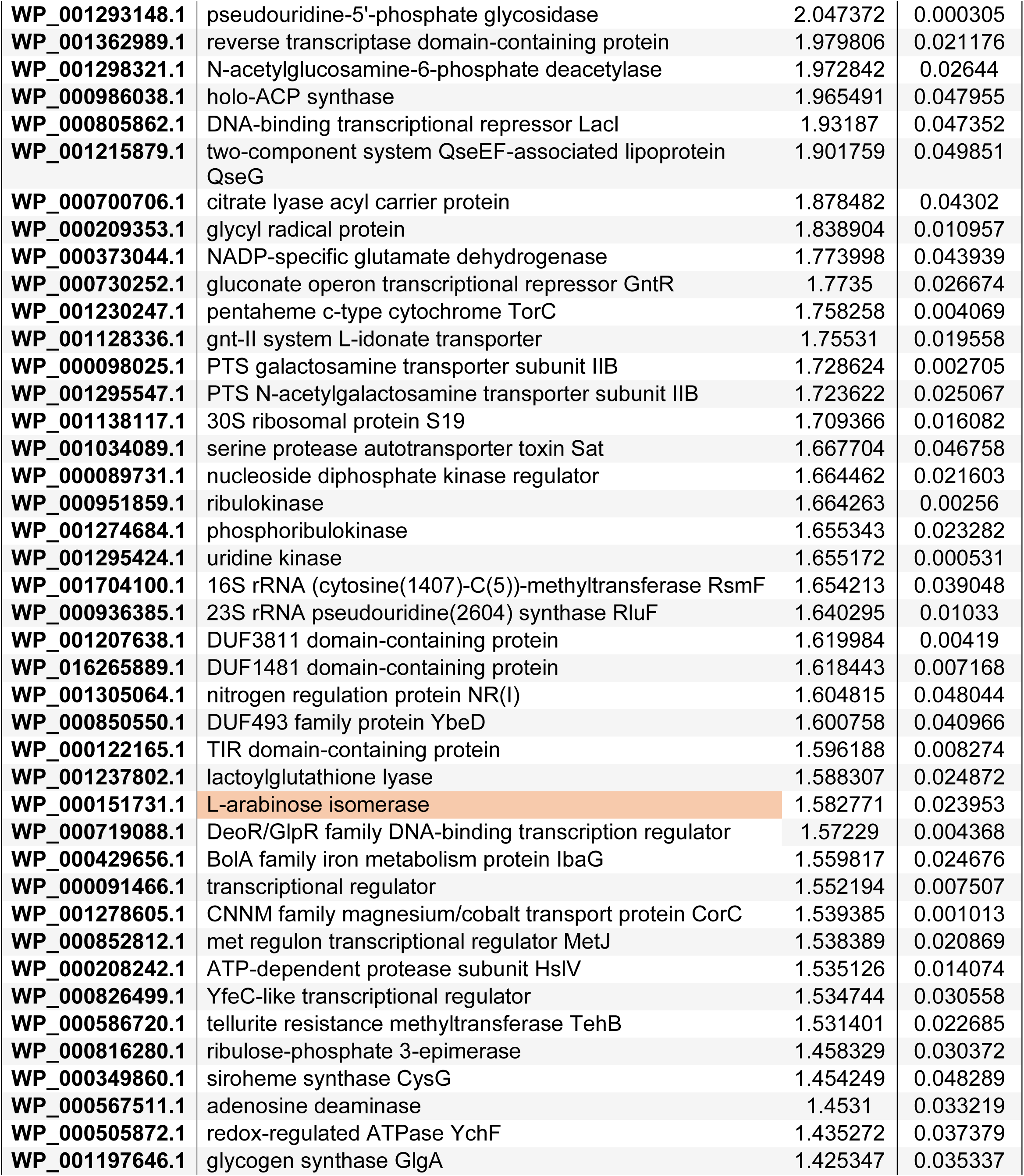
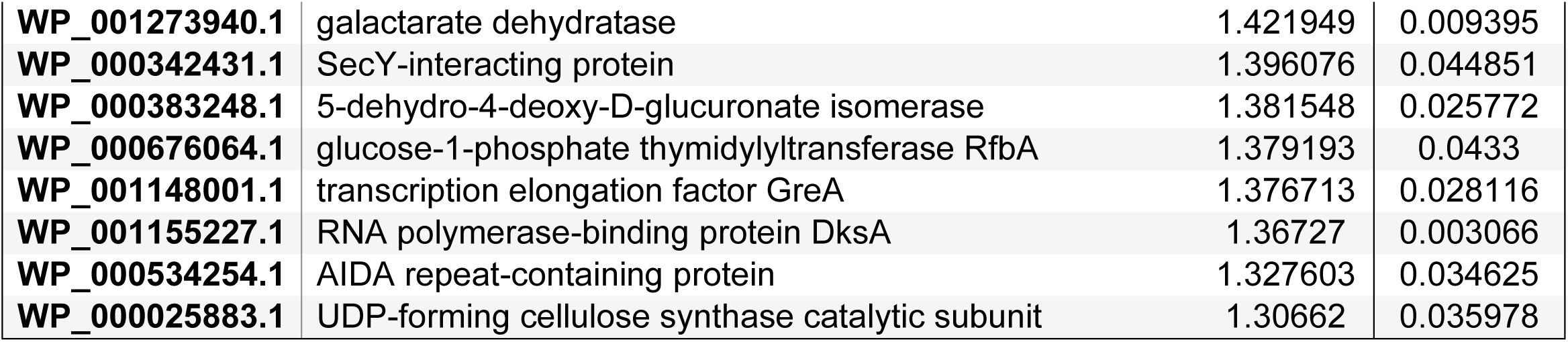
*E. coli* proteins enriched in triple species biofilms. Summary of proteins that specifically mapped to the *E. coli* CFT073 genome and were significantly enriched in triple species biofilms and not in dual-species biofilms.

