## Supplemental Figures and Table for "L-Fucose–Dependent Biofilm Formation by *Escherichia coli* Enhances Polymicrobial Interactions and Antibiotic Tolerance on Urinary Catheters"

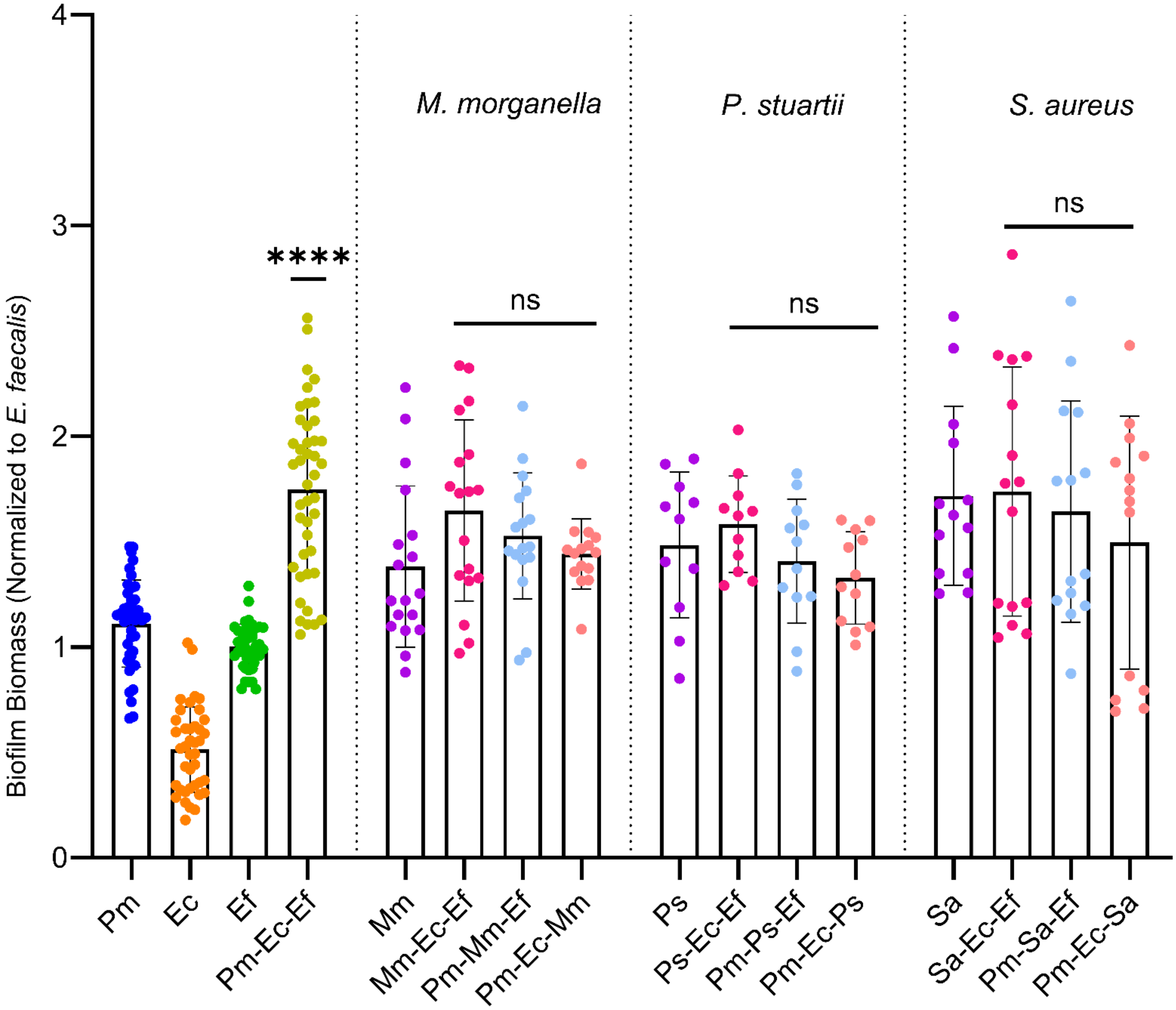


Supplemental Figure 1: Biomass of triple species biofilms formed with other common uropathogens. (A) Single and triple species biofilms were established in dilute urine for 24 hours and stained with crystal violet to determine biofilm biomass. Triple species biofilms are compared to the single species partner with the greatest biomass. Error bars represent mean and SDs from at least 4 independent experiments with 3 biological replicates. Statistical significance was determined by an ordinary one-way anova with multiple comparisons; *, P<0.05; ****, P<0.001; ns, non-significant.

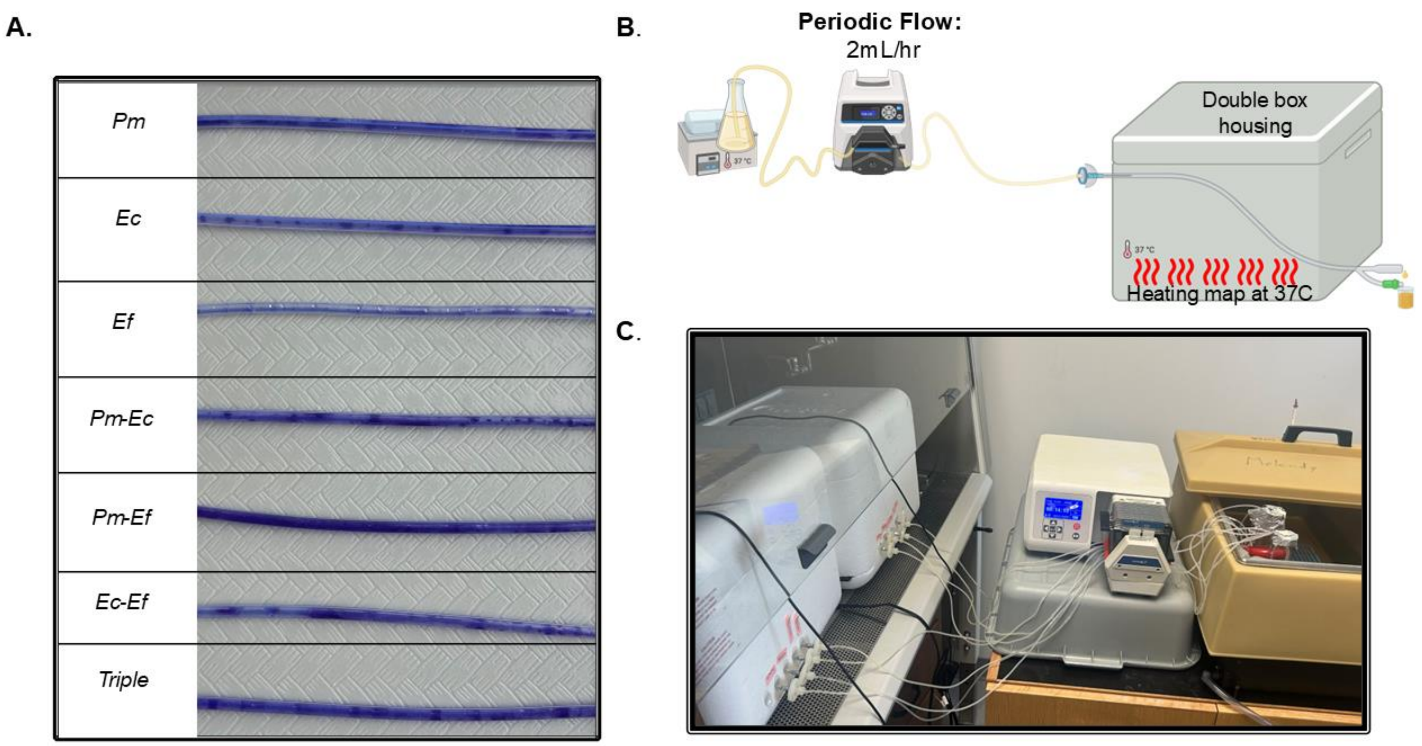


Supplemental Figure 2: Catheter biofilm reactor model. (A) Crystal violet stained catheters from 24 hours biofilms formed under flow in single, double, and triple species combinations. (B) Schematic of the catheter biofilm flow. (C) Image of biofilm reactor and flow set up.

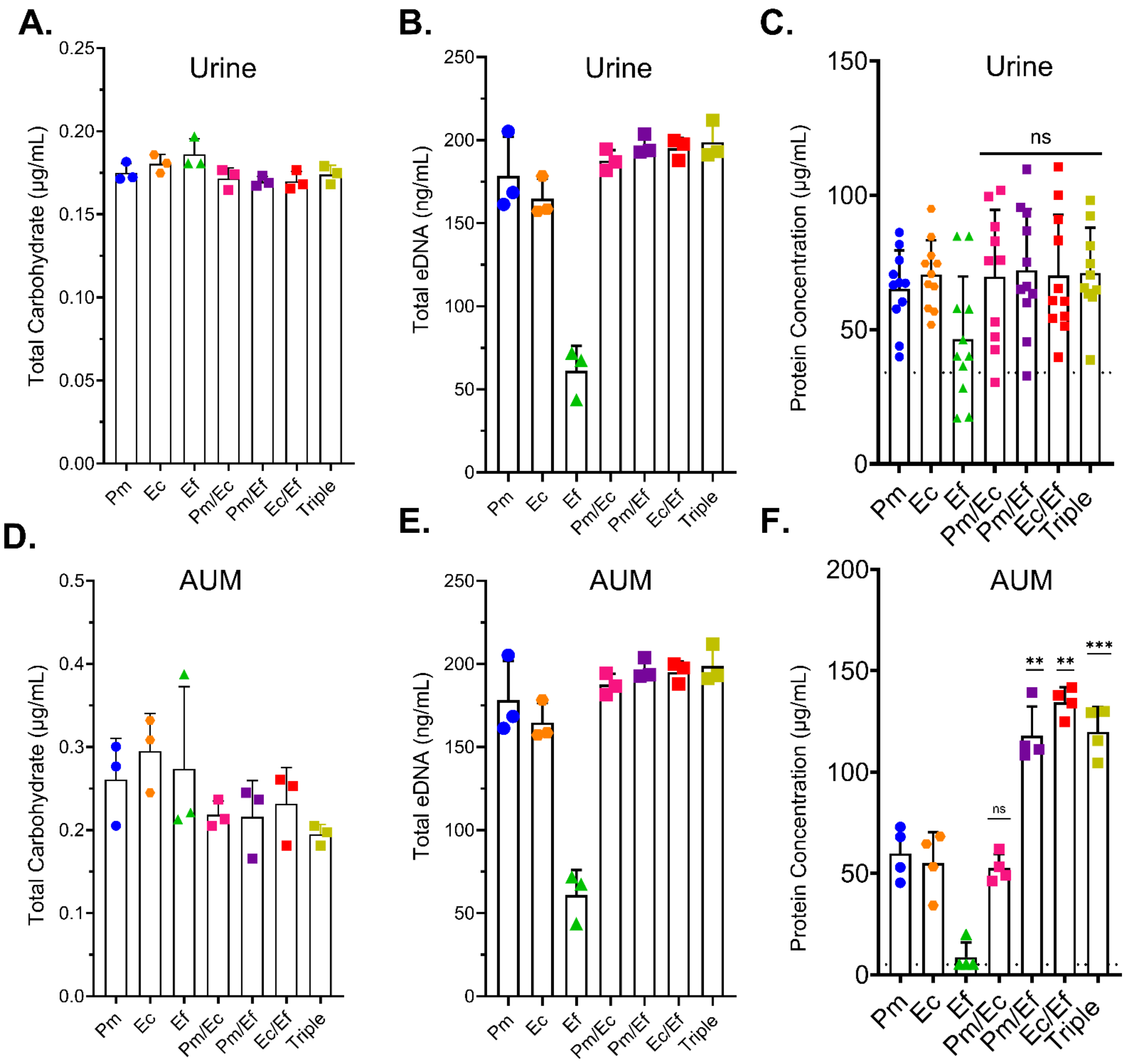


Supplemental Figure 3: Composition of single, double, and triple species biofilms formed in human urine and AUM. Single, double, and triple species biofilms were established in 24 well plates for 24 hours in undiluted human urine (A-C) or AUM (D-F) and composition was analyzed as follows: (A,D) total carbohydrate concentration, (B,E) total eDNA concentration, and (C,F) total protein concentration. Error bars represent means and SDs from at least 3 independent experiments. Statistical significance was determined by an ordinary one-way ANOVA test with multiple comparisons; *, P<0.05; **, <0.01; ***, P<0.001; ns, non-significant.

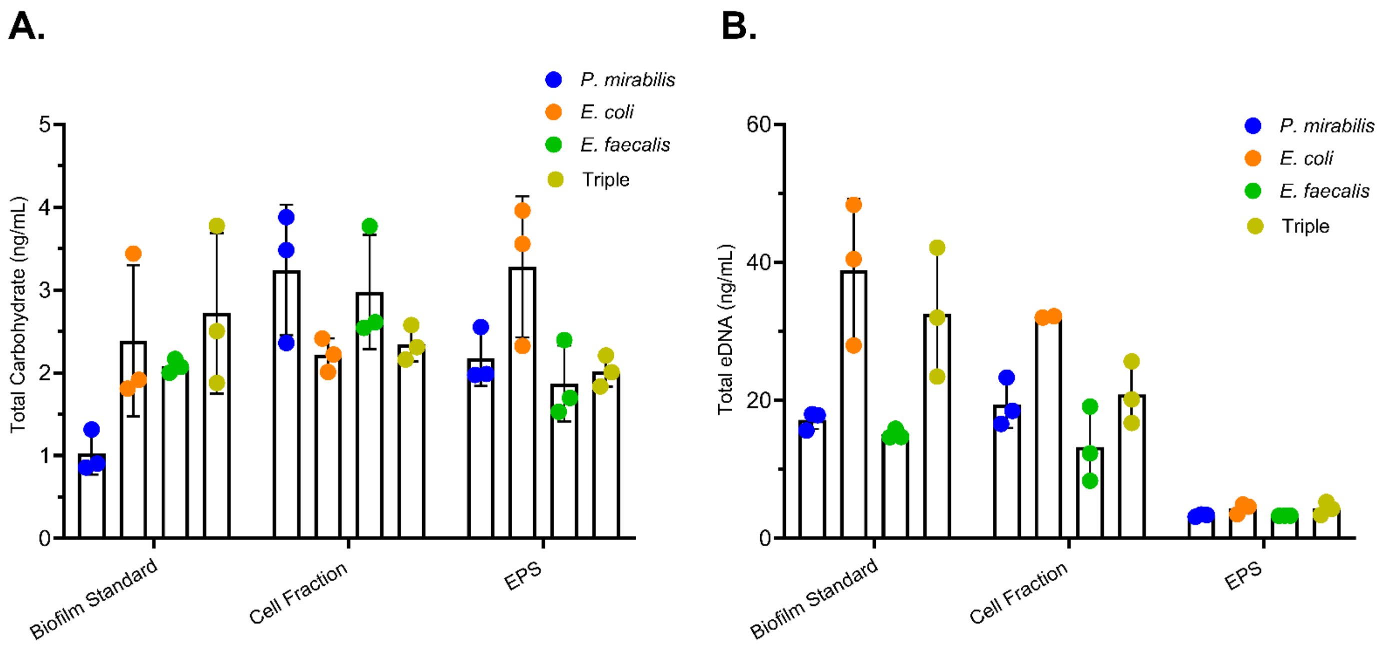


Supplemental Figure 4. Composition analysis of fractionated single and triple species biofilms. Single, double, and triple species biofilms were established in 24 well plates for 24 hours in diluted urine. Biofilms were separated into a cell-associated fraction and the EPS fraction and assessed for (A) total carbohydrate concentration and (B) total eDNA concentrations. Error bars represent means and SD from 3 independent experiments.

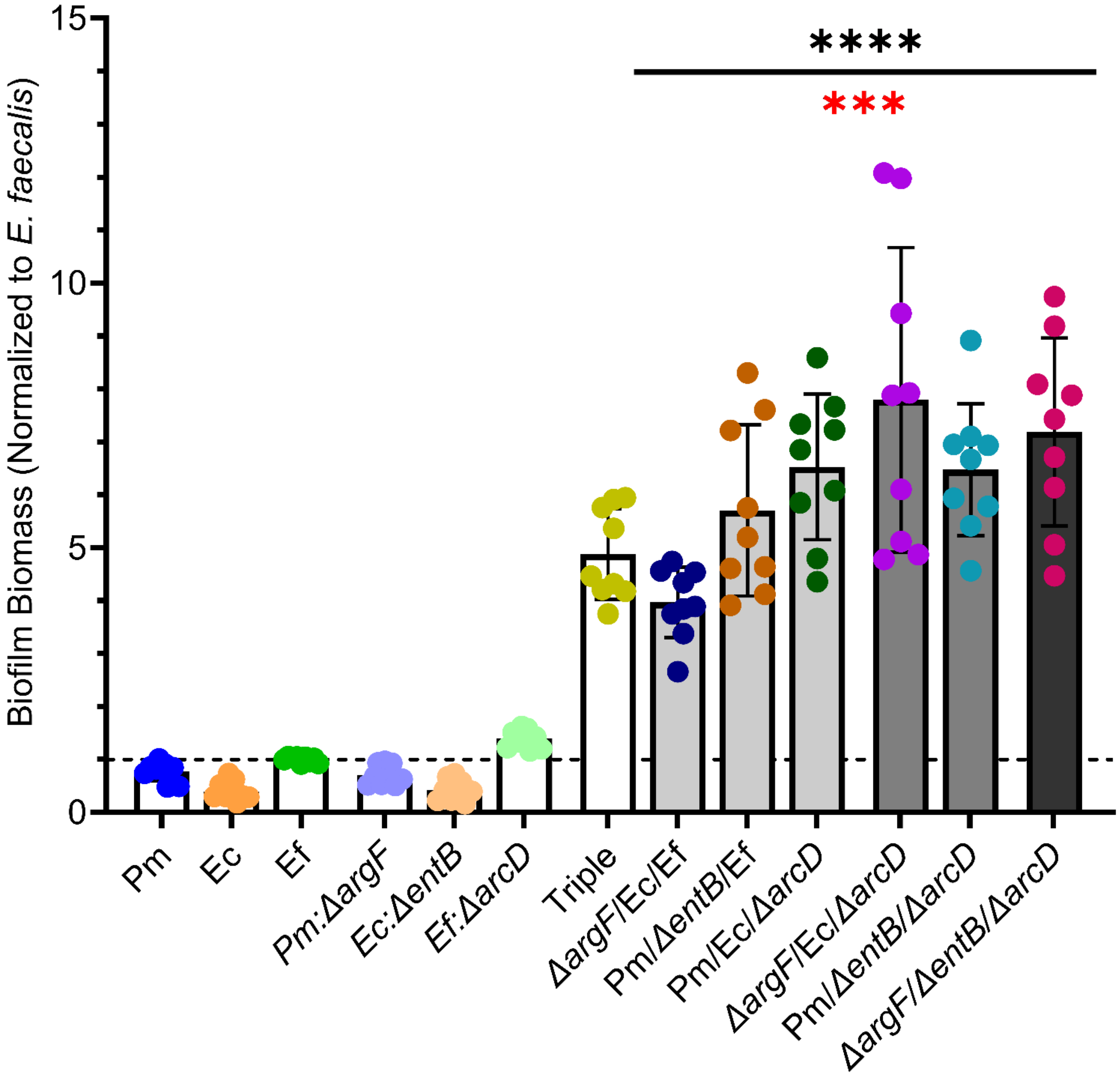


Supplemental Figure 5: Single, and triple species biofilm combinations with mutants, argF, entB, and/or arcD, involved in previously investigated mechanisms. Biofilm biomass was determined by crystal violet staining after 24 hours. Data points show mean and standard deviation from at least 3 independent experiments with 3 biological replicates each. Statistical significance was determined by a one-way ANOVA with multiple comparisons and compared to the single species *E. faecalis* (black, asterisk) and Ef:$\Delta arcD$ (red, asterisk); ***, P<0.001; ****, P<0.0001.

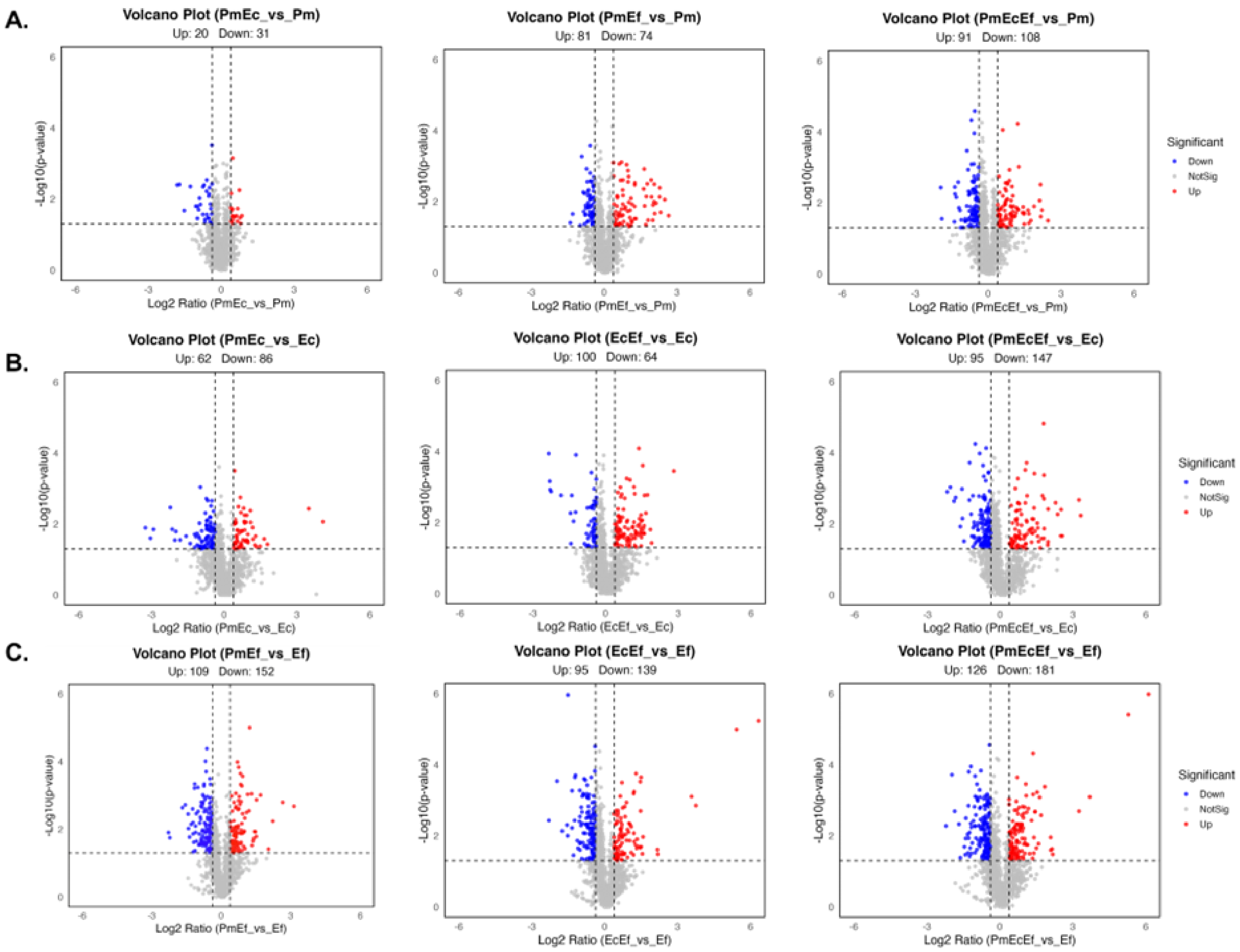


Supplemental Figure 6: Volcano plot analysis of differentially expressed proteins. Polymicrobial groups shown are compared to single species protein expression found in (A) *P. mirabilis,* (B) *E. coli,* or (C) *E. faecalis.* Enriched proteins (red) indicated have a fold change of >1.3 fold, depleted proteins (blue) have a <0.77-fold cutoff. Both groups have a statistical cut off of p=<0.05.

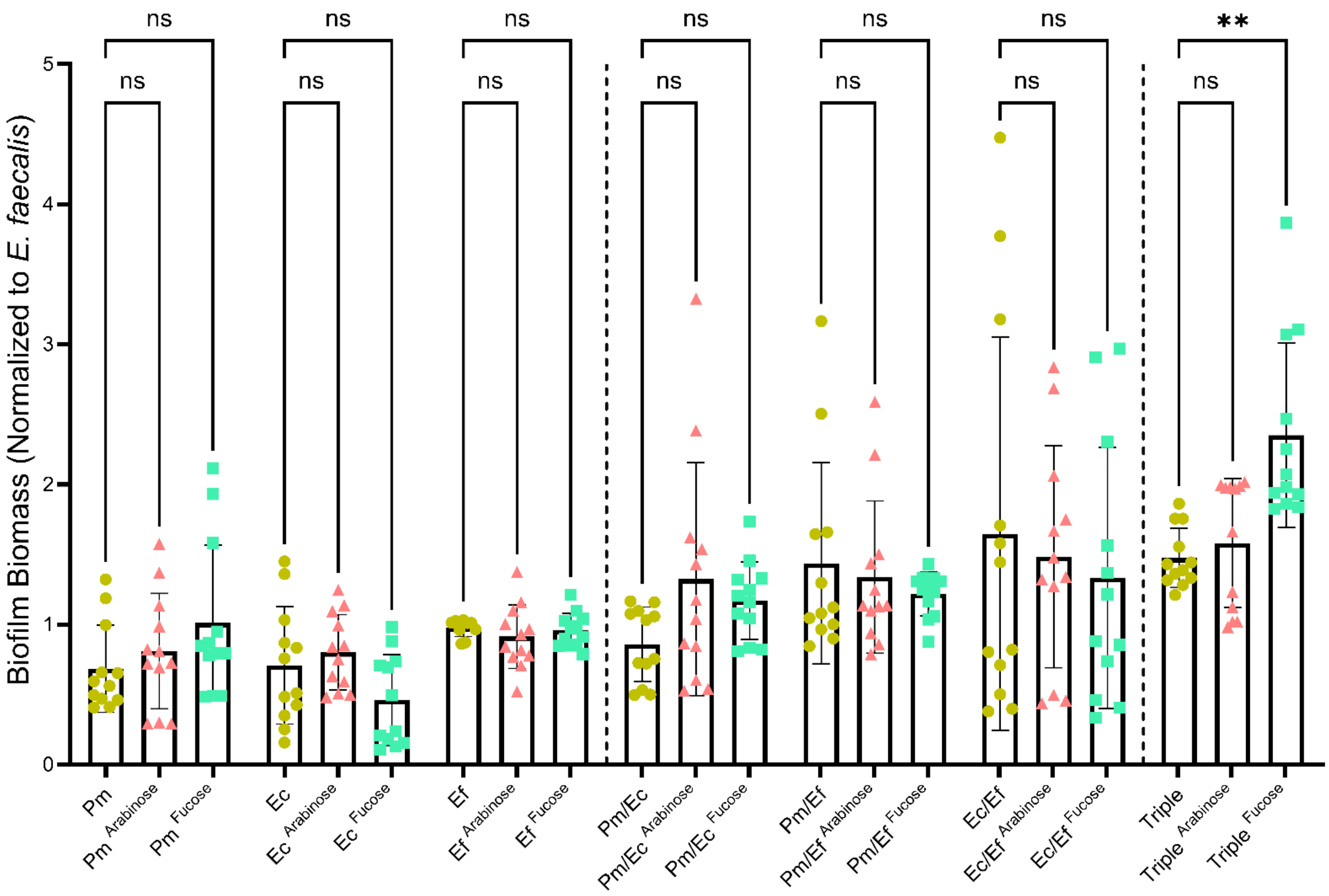


Supplemental Figure 7: Single, double, and triple species biofilms were established in 24 well plates in diluted human urine supplemented with 10 mM L-fucose or L-arabinose. Biofilm biomass was determined by crystal violet staining after 24 hours. Data points show mean and standard deviation from at least 4 independent experiments with 3 biological replicates each. Statistical significance was determined by a one-way ANOVA with multiple comparisons and compared to the untreated dilute urine groups; **, <0.01; ns, non-significant.
